# PiSpy: An Affordable, Accessible, and Flexible Imaging Platform for the Automated Observation of Organismal Biology and Behavior

**DOI:** 10.1101/2022.03.21.485129

**Authors:** Benjamin I. Morris, Marcy J. Kittredge, Bea Casey, Owen Meng, André Maia Chagas, Matt Lamparter, Thomas Thul, Gregory M. Pask

## Abstract

A great deal of understanding can be gleaned from direct observation of organismal growth, development, and behavior. However, direct observation can be time consuming and influence the organism through unintentional stimuli. Additionally, video capturing equipment can often be prohibitively expensive, difficult to modify to one’s specific needs, and may come with unnecessary features. Here, we describe the PiSpy, a low-cost, automated video acquisition platform that uses a Raspberry Pi computer and camera to record video or images at specified time intervals or when externally triggered. All settings and controls, such as programmable light cycling, are accessible to users with no programming experience through an easy-to-use graphical user interface. Importantly, the entire PiSpy system can be assembled for less than $100 using laser-cut and 3D-printed components. We demonstrate the broad applications and flexibility of the PiSpy across a range of model and non-model organisms. Designs, instructions, and code can be accessed through an online repository, where a global community of PiSpy users can also contribute their own unique customizations and help grow the community of open-source research solutions.

## Introduction

Observation remains a scientist’s most powerful and indispensable tool, especially in organismal biology. From the keen examinations of Jean-Henri Fabre to modern-day trail cameras, direct observation of organisms can yield both conclusions and new hypotheses [1]. However, observing key organismal behaviors may present challenges such as lengthy sessions, required nighttime monitoring, or unintentional and/or disruptive stimuli from the observer. Accessible and automated imaging equipment can ameliorate some of these challenges while still retaining the benefits of direct observation.

In recent years, increased efforts to develop and share open-source equipment for scientific research have provided accessible resources for both classical and modern experimental approaches. For example, there now exists affordable alternatives, commonly named “open labware,” for microscopy, optogenetics, centrifugation, and orbital shaking [2–5]. Open labware presents many key advantages over commercial alternatives. The reduced costs associated with open-source alternatives can benefit research labs with limited funding or allow for increased throughput with the use of numerous low-cost devices. These resources can also be used in STEM education where the high costs of specialized equipment may be prohibitive with typical teaching lab budgets. Additionally, open labware can be more customizable than commercially available versions, since their designs are distributed under open-source licenses, which generally allow users to alter designs at their will with appropriate credit to the original design [5]. Unlike proprietary versions that are protected under copyright licenses and cannot be altered, this enables researchers to design, build, and use equipment suited exactly to their needs rather than having to purchase alternatives that may come with unnecessary features or are not quite designed for the intended research purpose (summarized in [6]).

A key development for open labware development has been the rise of affordable single-board computers, the most popular of which is the Raspberry Pi [7]. Raspberry Pi computers have the functionality of a full computer at a modest cost and have been used to develop various open-source alternatives for scientific research. For instance, a Raspberry Pi, Arduino, and various 3D printed and commercial parts were used to build FlyPi, a microscope capable of light and fluorescent microscopy, as well as optogenetic and thermogenetic experiments [2]. Similarly, PiVR, a system to conduct closed-loop optogenetic experiments in freely-moving animals was developed using a Raspberry Pi [8]. Notably, these and other Raspberry Pi-based platforms come with the major advantages of open-source hardware; they are affordable, customizable, and accessible. Raspberry Pis can help create cheaper alternatives to preexisting equipment or can be used to develop equipment that is custom-made for specific tasks, especially given the compatibility of the Raspberry Pi with sensors and other additional features [7]. Raspberry Pi-based devices can be transformative and greatly decrease the cost of science in research and teaching environments. For example, FlyPi was introduced as a research tool to MSc students, PhD students, and senior faculty members at universities in sub-Saharan Africa. The flexibility of the FlyPi (and the Raspberry Pi itself) allowed it to be used both in its intended purpose as a microscope, but also as a medical diagnostic tool [2].

One area in which Raspberry Pis have been used is in automated video and image acquisition. Raspberry Pis can connect to a dedicated camera, and the picamera Python package allows for a high level of user control with regards to its settings and functionality [9]. Across a range of fields, various biological studies have used the Raspberry Pi as a camera device to monitor behavior, growth, and development of animals, plants, and bacteria [2,10–14]. For instance, the Raspberry Pi was used to create “POLIR,” an open-source, customizable system for high throughput analytical microbiology imaging, and “Picroscope” uses the Raspberry Pi for a microscope for simultaneous longitudinal imaging, with demonstrated applications in 2D and 3D cell/tissue cultures and whole organism imaging of frog and zebrafish development and planaria regeneration [10,14]. However, researchers are often required to program their Raspberry Pis and create setups specifically for their studies (e.g. [12]). While this can be an effective solution in labs that have both the required coding experience and development time to build a DIY research tool, it can be a significant barrier to other research groups. Therefore, we find there is a need for the development of an open-source camera device that is affordable, broadly applicable, and accessible to users with little or no programming experience.

We have developed PiSpy, a device that uses a Raspberry Pi, Pi Camera, and a combination of commercially available, laser cut, and 3D-printable parts. PiSpy can record high resolution images or videos, with an 8 MP camera that can record videos up to 1080p. After initial setup, PiSpy can operate without an active user to record at specified time intervals or when triggered by an external source, such as a motion sensor or IR break beam. Using custom-built LED light boards, we demonstrate that PiSpy can also automate a light-cycling program while also capturing behaviors throughout a 24-hour period. All functionality for PiSpy is controlled through an easy-to-use graphical user interface (GUI). PiSpy was designed to be affordable, easy to use, and flexible, and all code and designs have been posted to a Github repository (https://github.com/gpask/PiSpy) where users will be able to post their own modifications. A category on Gathering for Open Science Hardware (https://forum.openhardware.science/c/projects/pispy/58) has also been made to facilitate troubleshooting and other PiSpy related discussion. Our hope is that the PiSpy can be applied to a wide range of studies and contribute to the growing trend of powerful and accessible open-source research tools.

## Results

The base model of PiSpy for simple image or video capture is shown in Fig. 1A. A 3D printable holder mounts both the Raspberry Pi and Raspberry Pi Camera v2 to a laser cut wooden frame, allowing for easy height adjustment (Fig. 1A-B). The stands on the frame are also reversible, and a 3D-printed ball and socket mounts the camera, allowing for free rotation (Fig. 1C). Optionally, the Raspberry Pi computer can connect to input and/or output components, such as a motion sensor or lighting, respectively. This entire setup can be assembled for less than $100, or cheaper if multiple setups are being bought at once or certain features are omitted (S1 Table). A complete user’s guide and assembly manual can be found in the Supplementary Materials and will also be continually updated on the GitHub repository (https://github.com/gpask/PiSpy). The PiSpy GUI, written in Python3 using the TKinter package [15], controls its primary functionality. Features include capture mode, timed or input-triggered capture, light control, and camera resolution (Fig. 1D). For more advanced controls, such as changing the default image/video name and storage location or the specific camera settings, instructions are written in the user’s manual for how to edit these in the code itself.

**Fig. 1:**
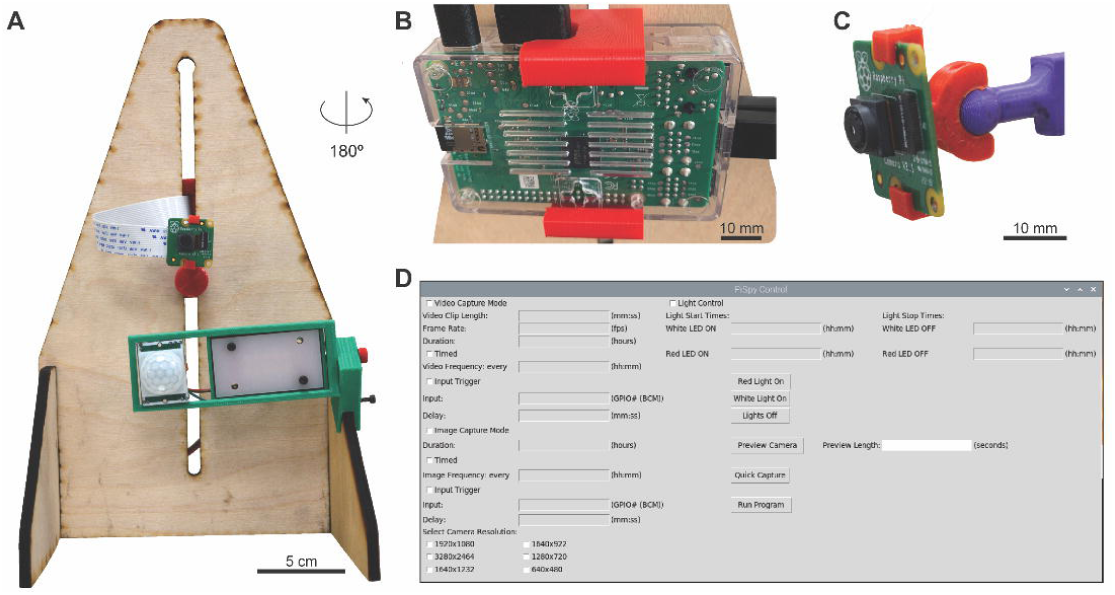
Overview of the PiSpy. **A**. Base model of the PiSpy, with optional LED lightboard and PIR motion sensor attached. **B**. Back of PiSpy highlighting 3D printed computer mount. **C**. 3D printed ball and socket camera joint, which allows for free rotation of the camera. **D**. Screenshot of the PiSpy GUI, which controls basic recording settings.

PiSpy can capture animal behavior at several scales (Fig. 2A-B). Imaging of *D. melanogaster* larval locomotion on an agar plate provided sufficient resolution for analysis with image processing software (Fig. 2A, S1 Video), and video of crayfish behavior when placed in an aquarium allowed for the observation of subtle movements of the swimmerets and legs, even though the animals were underwater and being viewed through a plastic container (Fig. 2B, S2 Video). Automated image capture also makes PiSpy an effective device for capturing and visualizing organismal growth over time (Fig. 2C-D). Time-lapse imaging of various beans (*Phaseolus vulgaris*) growing in clear planters showed detailed root and shoot growth (Fig. 2C, S3 Video). At a smaller scale, imaging of the soil bacterium *Bacillus mycoides* captured the growth and expansion process over time (Fig. 2D, S4 Video). These time lapses allow for clear visualization of organismal growth and could be used in a more formal study to compare and quantify growth of different species or under different growing conditions.

**Fig. 2:**
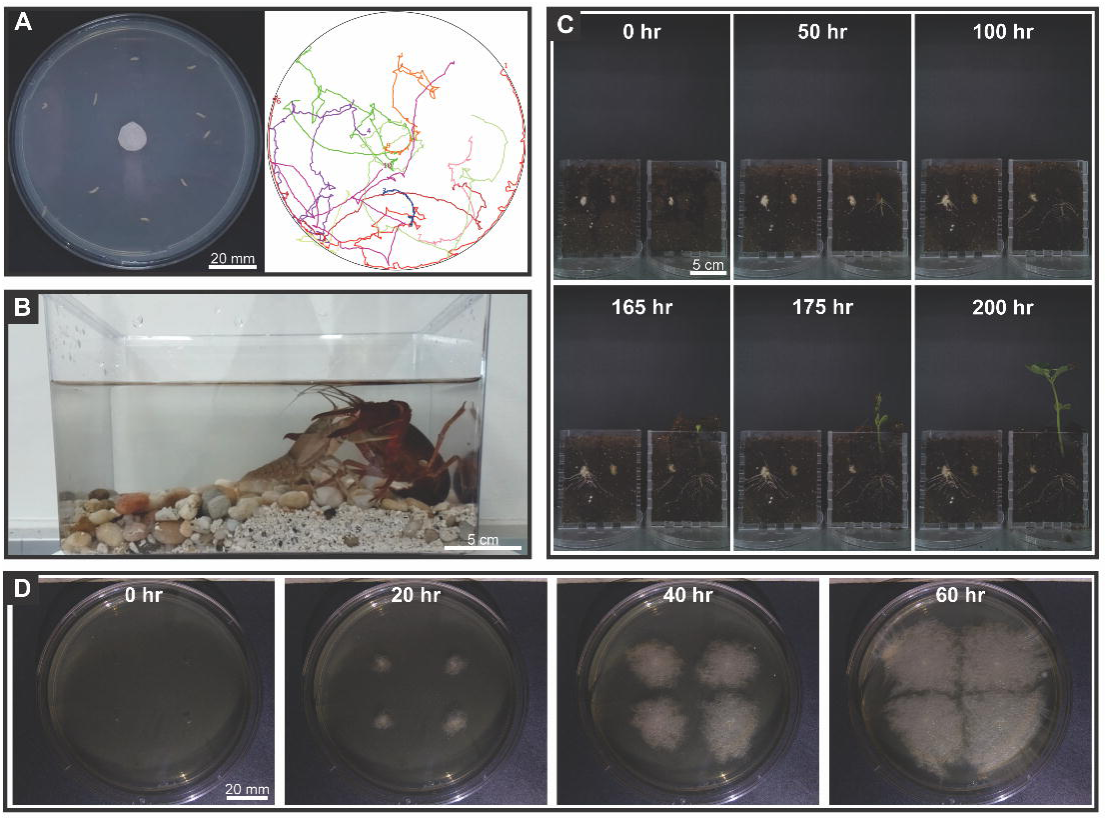
Video and time lapse image recording. **A**. Image of 10 *D. melanogaster* larvae initial position (left) and tracking data, with the number denoting that larva’s final position relative to central apple cider vinegar bait (right) (S1 Video). **B**. Image still of recording of crayfish dueling behavior. Two crayfish, one *Orconectes rusticus* and one *Procambarus clarkii* were placed in a container and the PiSpy was used to record their interaction for thirty seconds (S2 Video). **C**. Representative images from a time lapse video of *Phaseolus vulgaris* bean growing in clear planters and imaged by the PiSpy every 5 minutes (S3 Video). **D**. Representative images from a time lapse video (S4 Video) of *Bacillus mycoides* bacteria growing on a TSA+G agar plate and imaged by the PiSpy every 5 minutes (S4 Video).

PiSpy’s flexibility also allows it to be customized for more specific research purposes. For example, we have used PiSpy to monitor social behaviors in colonies of the Indian jumping ant, *Harpegnathos saltator*. Because the ants are housed in nestboxes of a fixed size, we have modified the wooden frame and mount to enclose the container to allow for easy overhead recording of the colony (Fig. 3A). To maintain a light-dark cycle for the ants, we used PiSpy’s LED light control capabilities to be able to record behaviors throughout the day. Custom LED printed circuit boards (PCBs) can be connected to the general-purpose input output (GPIO) pin of the Raspberry Pi and allow for the cycling of white and red lighting (Fig. 3A). In our experiment, the red light is not detected by the ants but allows for both day and night imaging (Fig. 3B-C, S5 Video, S6 Video). Specific camera settings are used to record in each different lighting conditions to ensure the desirable imaging quality. We have programmed in default camera settings for day and night recording, but the user’s manual provides instructions for how to modify these within the PiSpy code. For monitoring of organisms that would be affected by red light, infrared (IR) lighting and the Raspberry Pi NoIR camera could similarly be used to record at night. It is our hope that users will create their own modifications of the PiSpy hardware and/or software and will share these for use by other researchers and inspire further customizations.

**Fig. 3:**
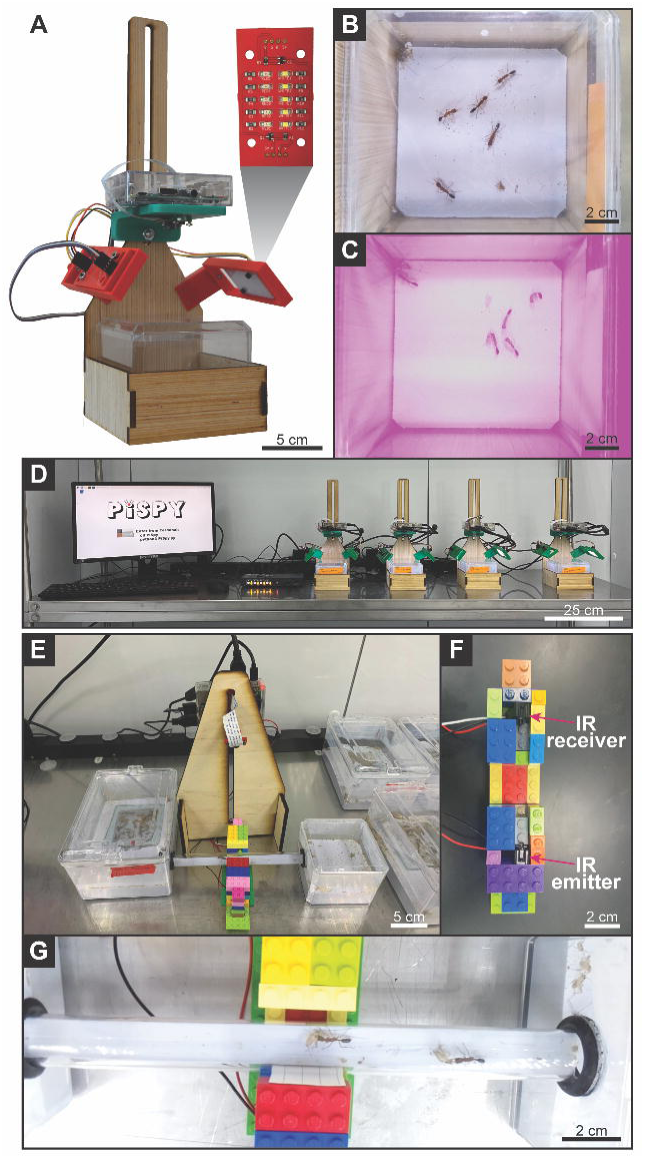
Monitoring *Harpegnathos saltator* ant behavior. **A**. Custom designed PiSpy setup for ant behavioral recording. The inset PCB boards provide LED lighting that cycles between red and white as specified by the user. **B**. Sample image from a video (S5 Video) of an *H. saltator* colony recorded during the day with white lighting from the PiSpy LED PCBs. **C**. Sample image from a video (S6 Video) of an *H. saltator* colony recorded during the night, with red lighting from the PiSpy LED PCBs. **D**. Multi-PiSpy setup using a KVM switch to control all four PiSpy units with a single monitor, keyboard, and mouse. **E**. Sample image still from the recording of *H. saltator* feeding. An IR break beam connected to the PiSpy was placed such that it would be triggered when an ant crossed a tube connecting a nest box to a foraging arena, and the PiSpy was used to record ants going to the arena and returning carrying crickets (S7 Video).

The GPIO pins of the Raspberry Pi can also be used to trigger image or video capture with an external sensor, such as a motion sensor or IR break beam. In our setup, an ant disrupts the IR beam as it walks to a foraging arena, and again when it returns to the main colony carrying a cricket (Fig. 3C, S7 Video, S1 Fig.). As an alternative to automated recordings at fixed time intervals, triggered recordings such as these could be used to monitor specific activities, such as feeding patterns or other behaviors.

## Discussion

Considering the current success of custom-developed, Raspberry Pi recording rigs in specific research purposes [12,13], there is a clear need for a broadly applicable, open-source, and accessible device that can be used in future studies. We have developed PiSpy to provide this resource while maintaining a low assembly cost and user-friendly interface. We hope that the open-source design allows for customization of both the hardware and software by researchers across the globe, and any updates or modifications made to PiSpy can be publicly shared on the GitHub repository and discussed on the GOSH forum. As demonstrated above, PiSpy can currently already be applied to a broad range of studies with various organisms, and its flexibility and ease of modification means it could easily be used in an immeasurable number of future experiments. Just as PiSpy was initially inspired by FlyPi, we hope scientists can use this as a springboard for their own innovations and ideas for open-source hardware in the future.

PiSpy can be especially useful in educational environments. Its low cost and ease of assembly mean it could easily be purchased in bulk (further reducing the individual cost, see Table S1), allowing for individual students or groups of students to each work with their own device. Additionally, because PiSpy can facilitate a broad range of research questions, students could use it to design their own independent experiments. Already, PiSpy has been used for undergraduate courses in biostatistics and entomology for independent projects as part of a course-based undergraduate research experience (CURE). PiSpy’s affordability and accessibility lends itself to informal learning environments, such as part of community science projects or curiosity-driven self-learning. Finally, while little to no experience in 3D fabrication, engineering, and coding is required to operate PiSpy, it presents the opportunity for motivated users to develop proficiency in these valuable skills.

PiSpy’s affordable, flexible, and accessible nature make it a robust tool for studying organismal growth, animal behavior, and a myriad of other possible applications. Its low cost and automated recording can increase throughput by using multiple devices, especially when coupled to powerful video analysis tools currently available [16]. As novel customizations are developed and shared online, we envision the evolution of the PiSpy as an emerging and increasingly functional research and teaching tool across the biological sciences.

## Methods

A complete User’s Guide and Assembly Manual is deposited on Github (https://github.com/gpask/PiSpy) and also included in the Supplementary Materials. For troubleshooting and other discussion, please use the forum on GOSH: https://forum.openhardware.science/c/projects/pispy/58.

### GUI

The PiSpy GUI was written in Python3 using the TKinter package to interface with the Tk GUI toolkit. It uses the GPAC MP4Box multimedia packager to convert .h264 files to .mp4. Different aspects of the PiSpy’s functionality are contained in their own classes, allowing for easier creation/alteration of functionality by users.

#### Bacteria isolation and growth

A small amount of soil from Middlebury, VT was placed in Trypticase Soy Agar+Glucose (TSA+G) broth containing 17 g/L tryptone, 3 g/L soytone, 10 g/L dextrose, 5 g/L NaCl, 2.5 g/L K_2_HPO_4_, and 15 g/L Agar. The broth was placed in an 80°C water bath for 10 minutes, then removed and allowed to incubate at 25°C for 30 minutes. Using a sterile loop, the broth was streaked onto a TSA+G plate and allowed to incubate overnight at 25°C. One large colony was selected from the plate and used to inoculate a separate TSA+G plate in four places. Using the PiSpy, images were recorded every 5 minutes for 95 hours, and QuickTime Player on Mac was used to create a time lapse video at 29.97 frames per second from the ensuing image stack.

#### Bean growth and imaging

Two clear 5.15’ x 4.38” x 1.5” plexiglass planters (fabricated using a laser cutter) were filled with Miracle Gro® indoor potting mix soil, and two varieties of the common bean *Phaseolus vulgaris* were planted on the edge of each planter such that they were visible from one side. The design of the relatively thin planters optimized root visibility, but other options can also work. The planters were watered immediately after the beans were planted, and again whenever the soil appeared visibly dry. Using the PiSpy, images were recorded every 5 minutes for 11 days, and QuickTime Player on Mac was used to create a time lapse video at 29.97 frames per second from the ensuing image stack.

#### Drosophila larvae locomotion

10 third instar *D. melanogaster* larvae were placed on an agar plate. A filter paper disc was placed in the center and undiluted apple cider vinegar was applied onto the disc to attract the larvae. Using the PiSpy, a 5-minute video was recorded. Subsequently, the MTrackJ plugin for ImageJ was used to analyze the movement of the larvae by manually annotating the location of the anterior side of the larvae in every frame of an image stack generated by taking 1 frame from each second of the video[17].

#### Crayfish observation

Two crayfish, one *Orconectes rusticus* and one *Procambarus clarkii* were placed together in a clear plastic container. The PiSpy was used to record a video of their interaction for 30 seconds.

#### Ant observation

Five *Harpegnathos saltator* ants were isolated from an established colony and placed in a new nest box. The PiSpy was used to record 30 second videos every 20 minutes for 3 days.

For the motion triggered setup, a large ant nest box was connected to a smaller foraging arena by a tube. An IR break beam motion sensor connected to the PiSpy and held secure by a custom-built LEGO® structure was placed such that it would be triggered when an ant passed through the connecting tube. Crickets were placed in the foraging arena, and the PiSpy was set to input trigger mode, recording 15 second videos when the break beam was triggered with a delay of 5 seconds after video acquisition.

## Supporting information

PiSpy User's Guide

## Author Contributions

BIM: Conceptualization, Software, Formal Analysis, Investigation, Methodology, Resources, Supervision, Validation, Visualization, Writing – Original Draft Preparation, Writing – Review & Editing

MJK: Conceptualization, Software, Formal Analysis, Investigation, Methodology, Resources, Supervision, Validation, Visualization

BC: Conceptualization, Software, Investigation, Validation

OM: Methodology, Resources

AMC: Supervision, Validation, Writing – Review & Editing

ML: Conceptualization, Methodology, Resources, Supervision

TT: Conceptualization, Methodology, Resources

GMP: Conceptualization, Funding Acquisition, Investigation, Methodology, Project Administration, Resources, Supervision, Validation, Visualization, Writing – Review & Editing

## Acknowledgements

We would like to thank Tim Allen, Ciara Burke, Amanda Crocker, Erin Eggleston, Eric Moody, and Megan Warner-Hough for assistance in providing organisms and supplies for observation. We also thank Daphné Halley, Aiden Masters, Anthony Marinello, Sophia Marlino, Jen McGann, Andrew Ng, and Joanna Raup-Collado for testing and assembly troubleshooting efforts, Andrew Hawk for initial GUI development in guizero, and Joanna Pask and Elliot Pask for lending beans and LEGO® bricks. And a special thanks to Eamon McMahon for fabrication of experimental apparatuses and Andrea Vaccari for assistance in GUI coding. We would also like to thank the Maker-E makerspace at Bucknell University, the Middlebury College Makerspace for fabrication equipment and supplies, and the Biomedical Engineering and Electrical and Computer Engineering Departments at Bucknell University for fabrication support.

## Supporting information S1 Table. Bill of Materials

List, costs, and suggested links for all required parts to build the PiSpy, as well as optional additional equipment. For assembly and setup instructions, refer to the user’s guide (link once uploaded).

(XLSX)

**S1 Video. *Drosophila* larval tracking with MTrackJ on ImageJ (related to Fig. 2A)**.

(MOV)

**S2 Video. Crayfish dueling behavior (related to Fig. 2B)**.

(MOV)

**S3 Video. Timelapse video of *Phaseolus vulgaris* bean growth (related to Fig. 2C)**.

(MOV)

**S4 Video. Timelapse video of *Bacillus mycoides* colony growth (related to Fig. 2D)**.

(MOV)

**S5 Video. Daytime recording of *Harpegnathos saltator* behavior (related to Fig. 3B)**.

(MOV)

**S6 Video. Nighttime recording of *Harpegnathos saltator* behavior (related to Fig. 3C)**.

(MOV)

**S7 Video. Break beam triggered video of *Harpegnathos saltator* foraging behavior (related to Fig. 3D)**.

(MOV)

