## Supplementary material for "PiSpy: An Affordable, Accessible, and Flexible Imaging Platform for the Automated Observation of Organismal Biology and Behavior": PiSpy User's Guide

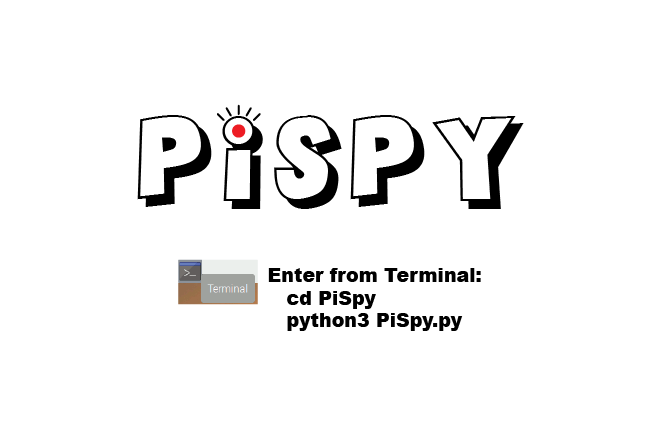

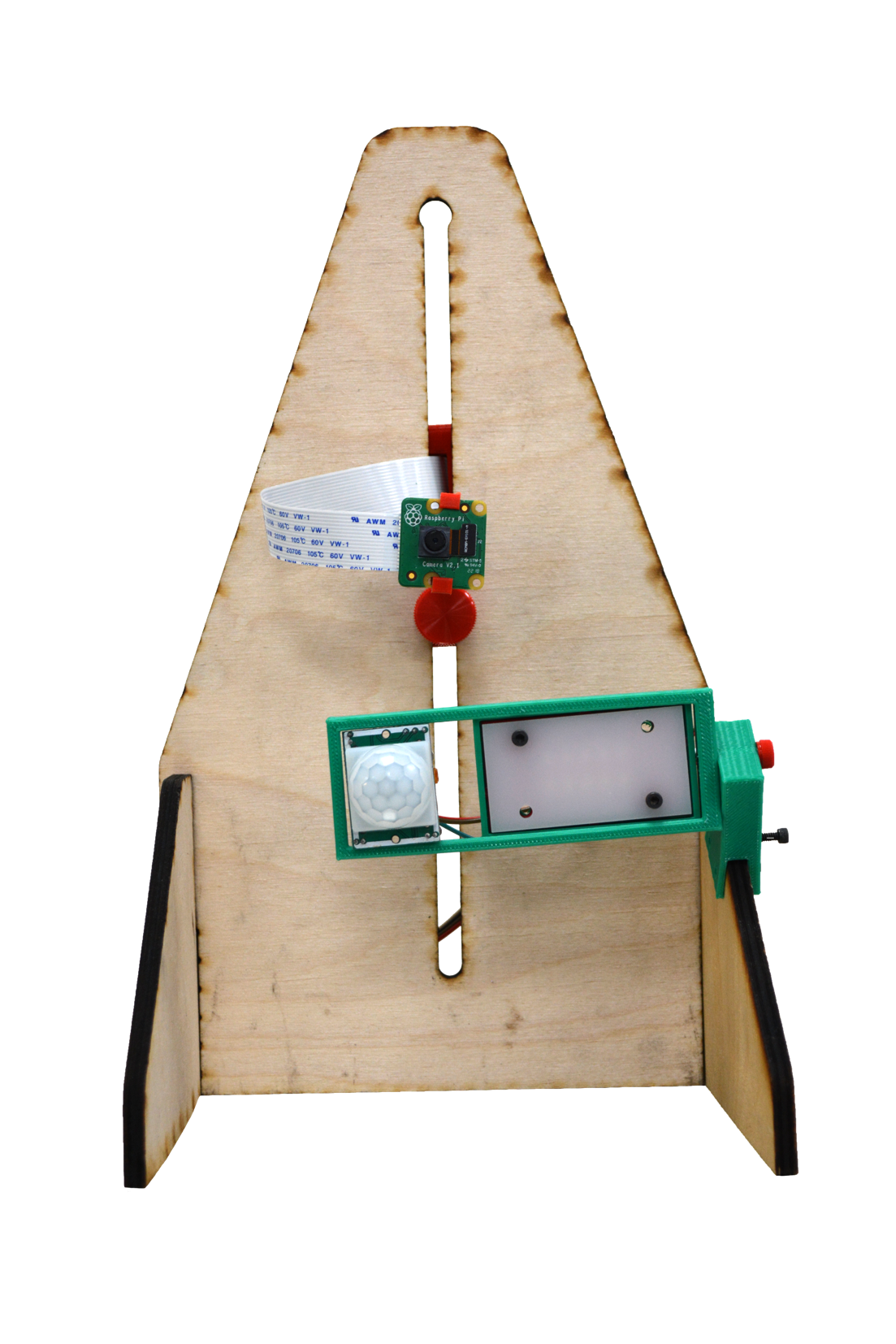

The PiSpy is an affordable device used to automate the acquisition of images and videos. It was first introduced on bioRxiv in 2022 (link). It uses a Raspberry Pi computer and camera to record video and images at specified time intervals or when triggered by an input signal, such as a motion sensor or force plate. Additionally, it can automate the control of LED lights. All settings for the PiSpy are controlled through a graphical user interface that uses the TKinter Python package. The following user guide will outline how to set up and operate the PiSpy. Our testing was with a Raspberry Pi3b or Pi4, which is primarily what this guide is written for, but with some modifications the PiSpy can also be used with older Raspberry Pi models.

### Setting up the Raspberry Pi

#### Installing Raspberry Pi OS

The first step to running the PiSpy is to set up the Raspberry Pi OS operating system on the computer. To do so, on a computer with an SD card reader, download the [Raspberry Pi imager](https://www.raspberrypi.org/software/). Insert the micro SD card into the computer and open the imager.

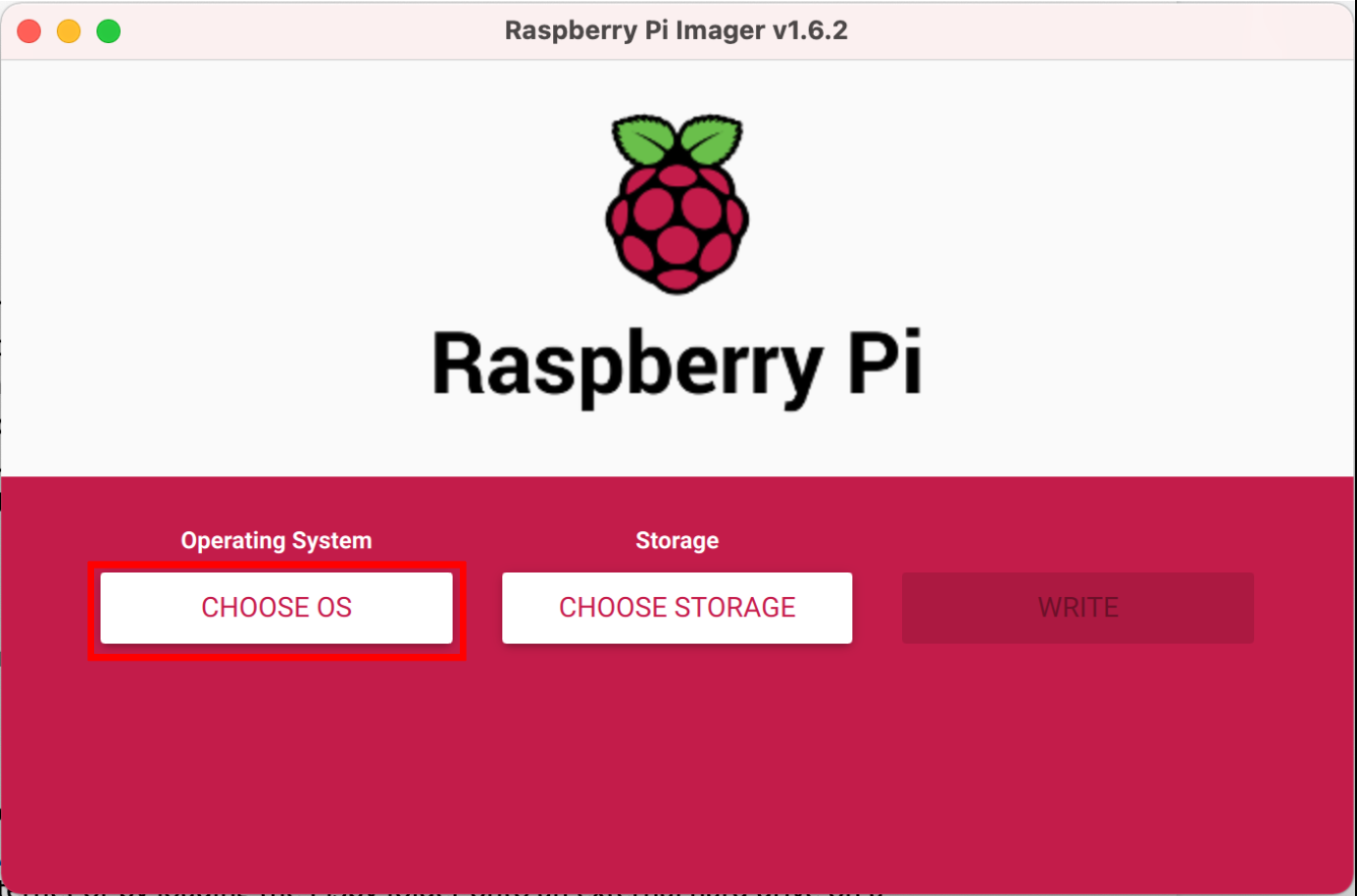

Select “CHOOSE OS,”

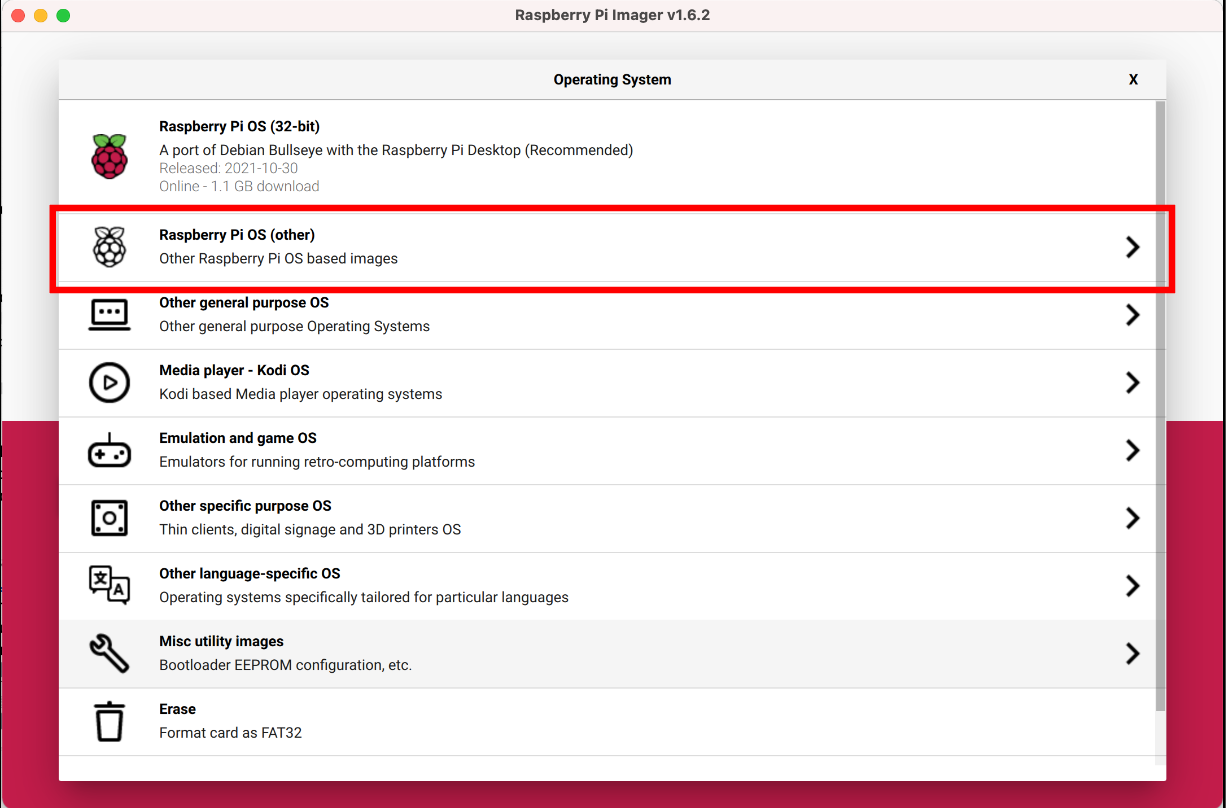

then select “Raspberry Pi OS (other),”

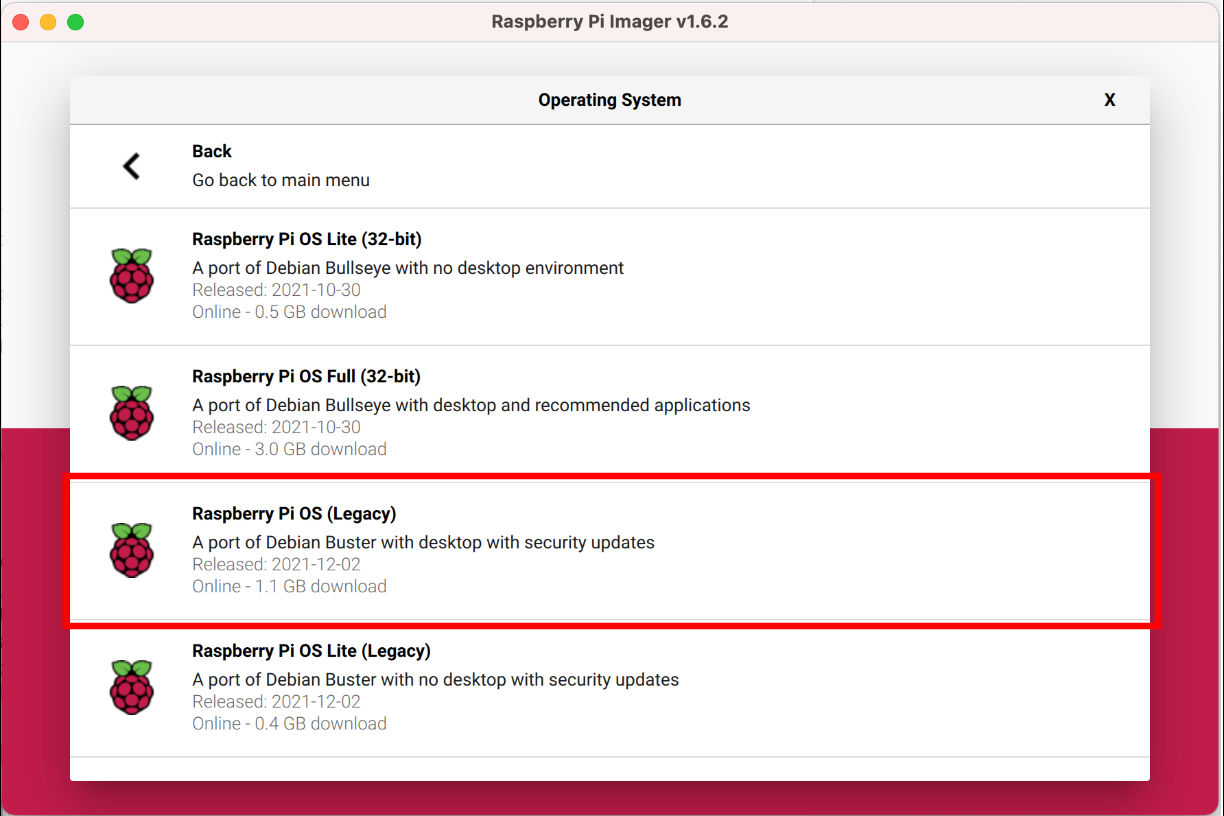

and select “Raspberry Pi OS (Legacy).”^[1]^

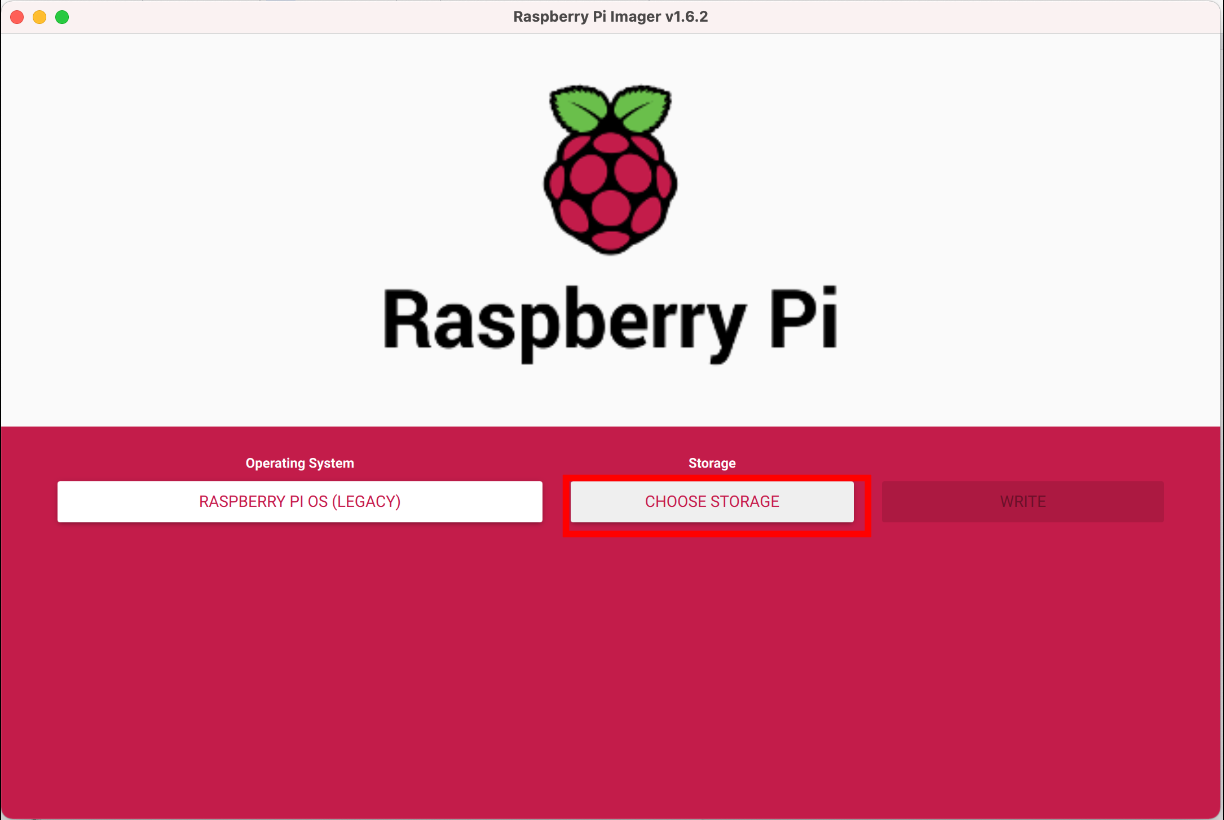

Next, select “CHOOSE STORAGE”

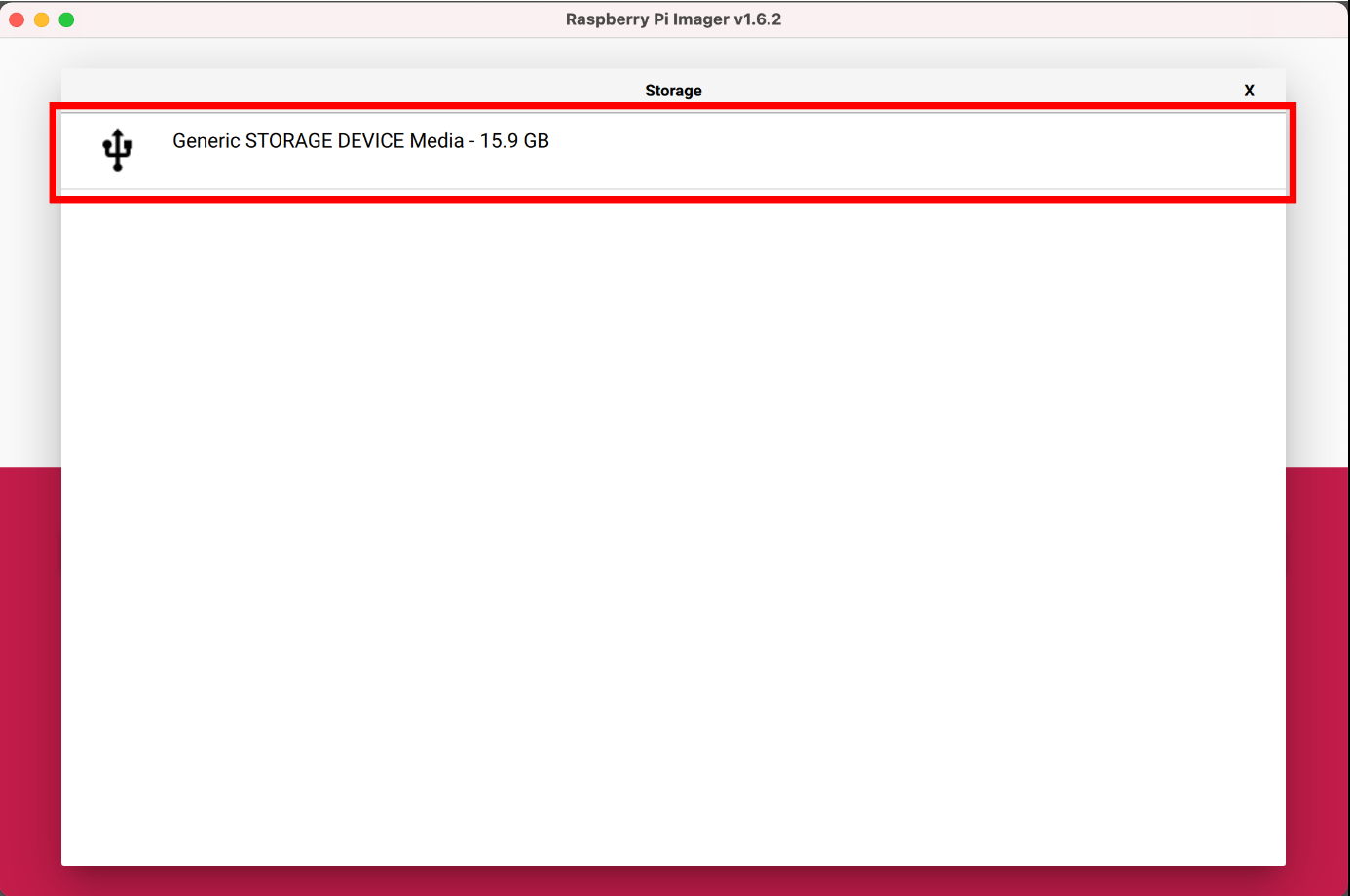

and select the SD card to be used for your PiSpy.

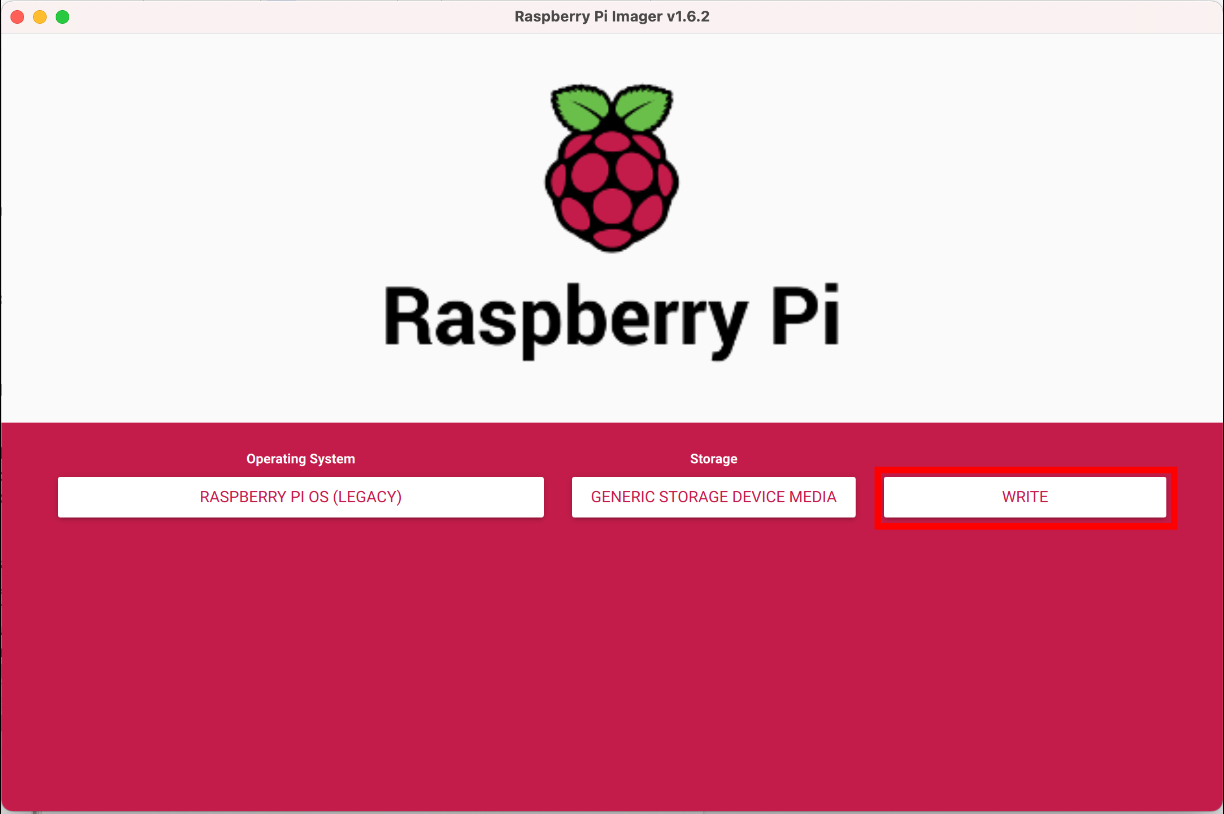

Finally, write the OS to the SD card by selecting “WRITE”

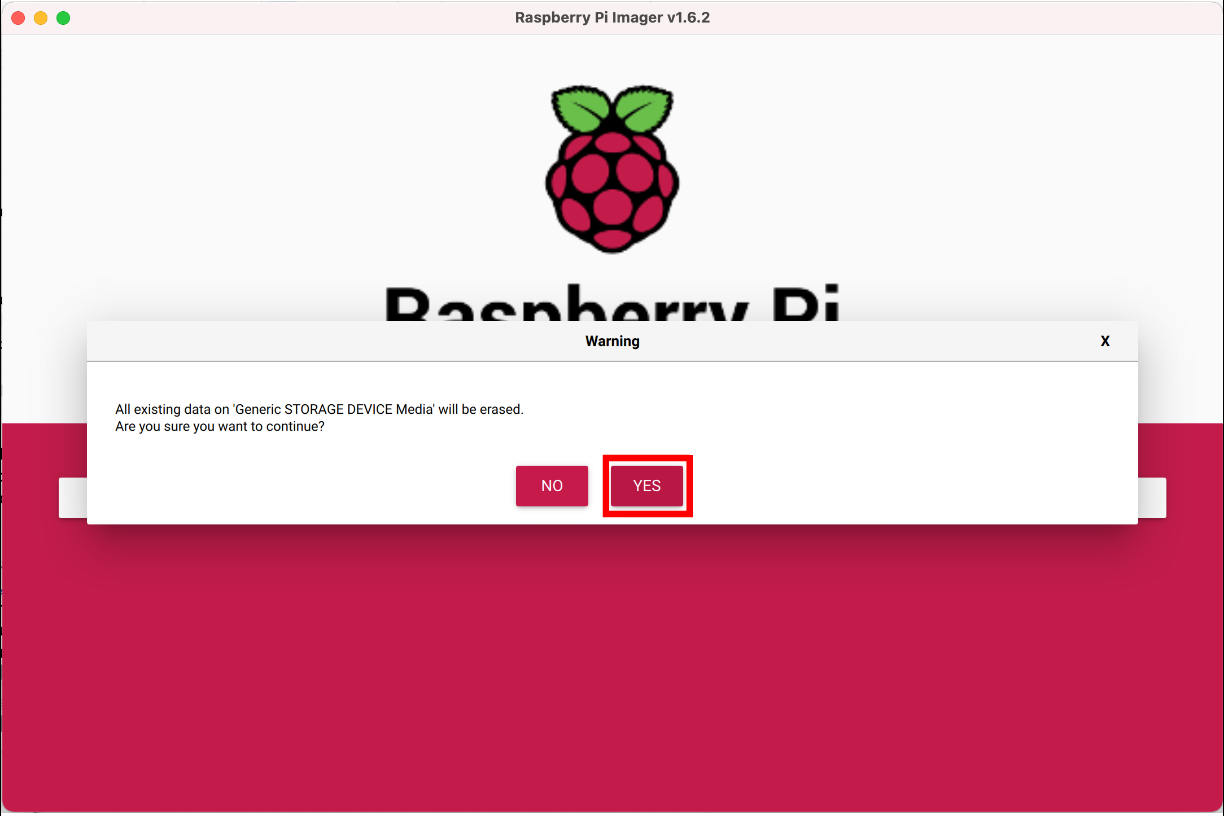

and confirming that this will erase any previous data on the SD card.

#### Installing the PiSpy code

For the graphical user interface (GUI), download the PiSpy folder from Software folder on the GitHub repository (<https://github.com/gpask/PiSpy/tree/master/Software/>). Upload it to a USB flash drive, then transfer the PiSpy folder to the /home/pi folder on the Raspberry Pi (see image). Alternatively if the Raspberry Pi has an internet connection, download the folder directly onto the Pi and move it from downloads to the /home/pi/ folder:

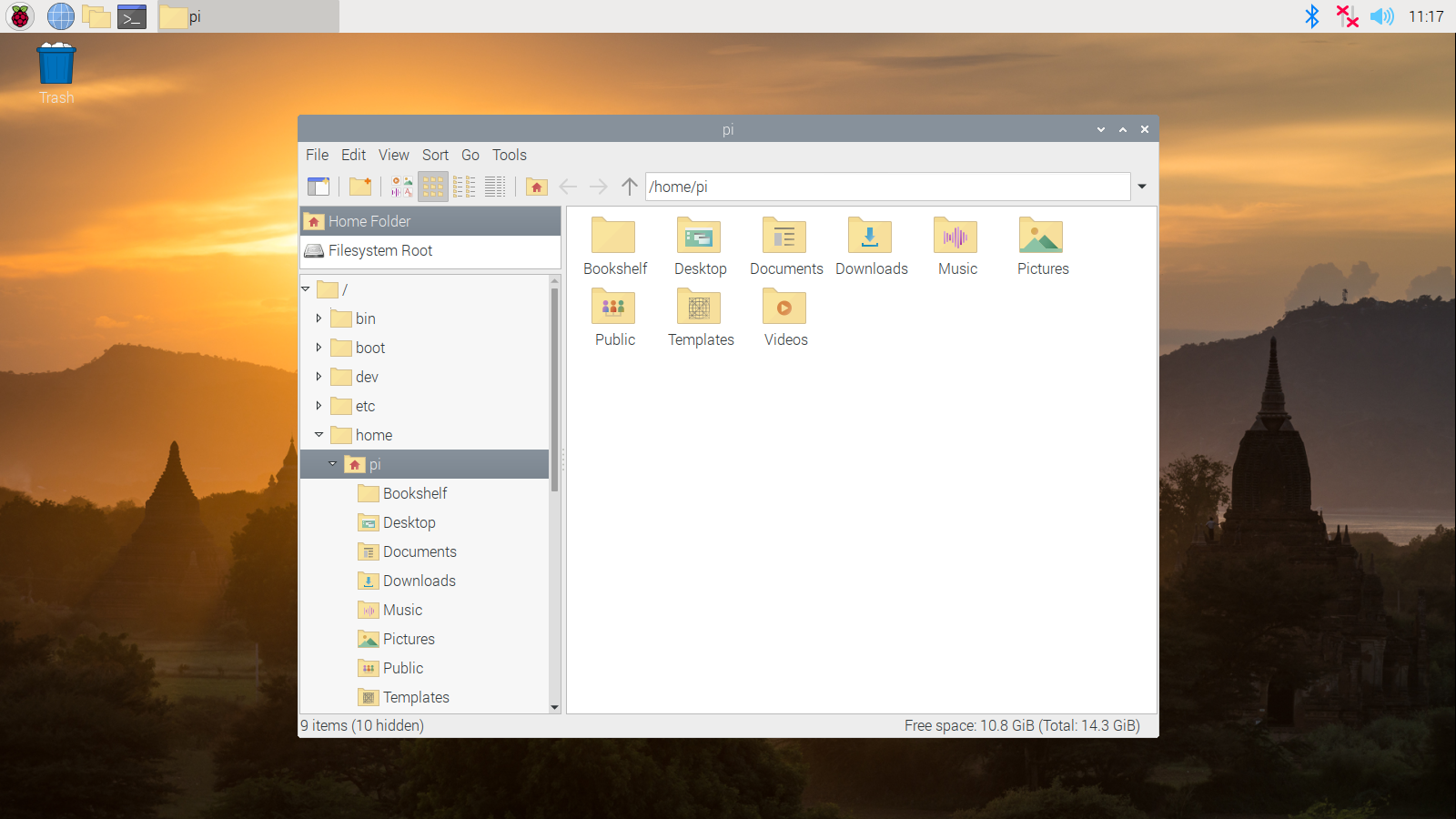

#### Enabling the camera

The first time the PiSpy is used, the user will also need to connect and enable the PiCamera by following the instructions here: <https://projects.raspberrypi.org/en/projects/getting-started-with-picamera/2>. If the connection between the camera and the Raspberry Pi is not secure, the PiSpy will give the following error message when you try to record an image: “picamera.exe.PiCameraMMALERROR: Failed to enable connection: Out of resources”. If this happens, reconnect the camera and make sure the connection is secure before enabling it on the Raspberry Pi, then restart the computer.

#### Installing GPAC

Additionally, upon first setting up the PiSpy, the user will need to install GPAC, which contains the MP4Box software the PiSpy uses to convert *.h264 video files to *.mp4. To do so, on a Raspberry Pi that is connected to the internet, type “sudo apt-get update” to update the system, then “sudo apt-get install -y gpac” to install the package. If your Raspberry Pi does not connect to the internet, GPAC cannot be installed. Comment out (place a # before) line 35 in the Record.py file. This will result in videos being saved as .h264 files, which can be converted to .MP4 files on another computer.

### Operating the graphical user interface

To access the GUI, open the command line by clicking on the terminal icon on the top bar of the screen. Then, enter the following commands, first to move to the PiSpy folder and then to open the PiSpy Control window, which is where the PiSpy is operated from:

cd PiSpy

python3 PiSpy.py

If you have already entered those commands, you can use the up arrow to access previously used commands in terminal. Also note that the cd PiSpy will only work if the folder containing the PiSpy code is still named “PiSpy.” If the name is changed for some reason (as can happen when unzipping the file), either change the name back to PiSpy or enter the new name for the folder when typing that command.

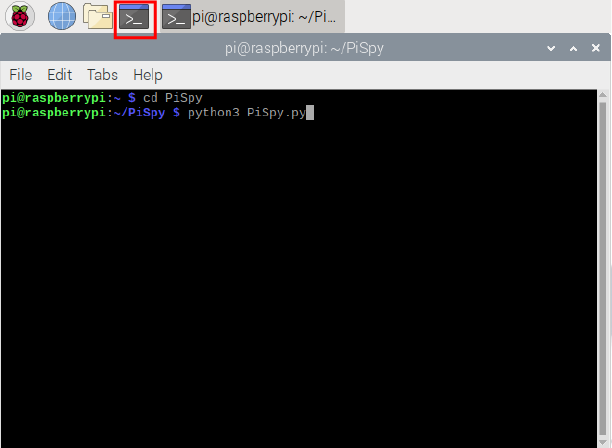

To help remember how to access the GUI, we have included on the Github repository a suggested background for the PiSpy (“PiSpy desktop.png”). To set this as the computer background, go to Preferences → Appearance Settings, then save the new image file to the PiSpy and select it as the desktop image.

Once the GUI is opened, choose your desired capture mode (image or video) and frequency (timed or triggered) by checking the corresponding boxes. Only one mode and frequency combination (i.e. timed videos) should be selected at a time. Additionally, select the camera resolution by checking the box of the desired resolution, and, for video recording, specify the frame rate. If no resolution is selected, the PiSpy will by default record with a resolution of 1920 x 1080. This link (<https://picamera.readthedocs.io/en/release-1.12/fov.html#camera-modes>) lists the possible resolutions and framerates for the V1 and V2 PiCameras. Depending on the videos being recorded and the quality of the microSD card used, selecting frame rate/resolution combinations at the upper limits of the accepted range can result in skipped frames. If this occurs, lower either the resolution or the frame rate.

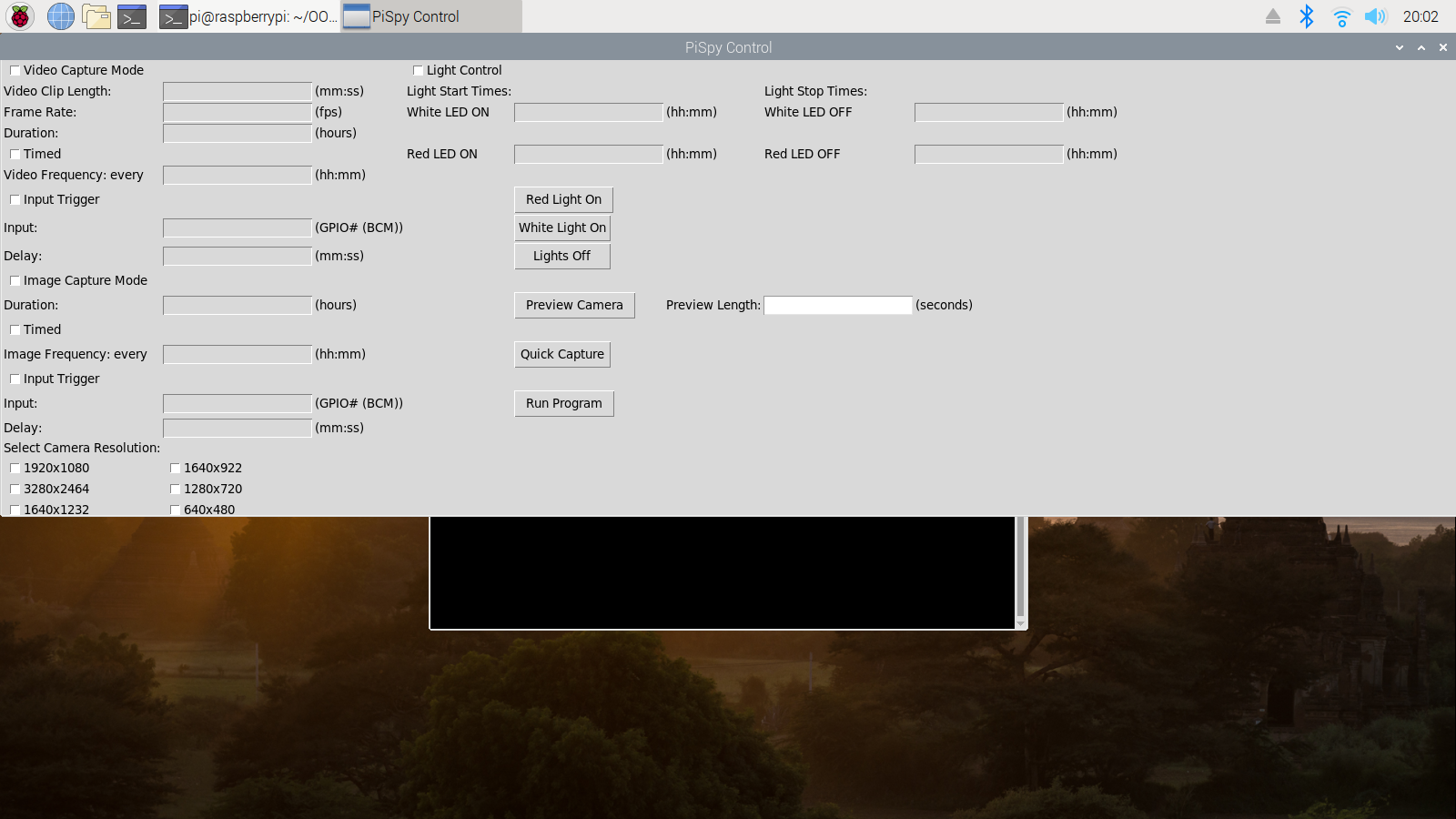

*Timed recording:*

Timed recording can be used when the user wants images or videos to be recorded at fixed intervals. The image or video frequency should be input in the hh:mm format (using a 24 hour clock), and specifies how often images/videos are recorded. For instance, in the below image, 30 second videos with a frame rate of 20 fps and a resolution of 1920x1080 will be recorded every 1.5 hours for 3 days. Note that the frequency must be less than 24 hours.

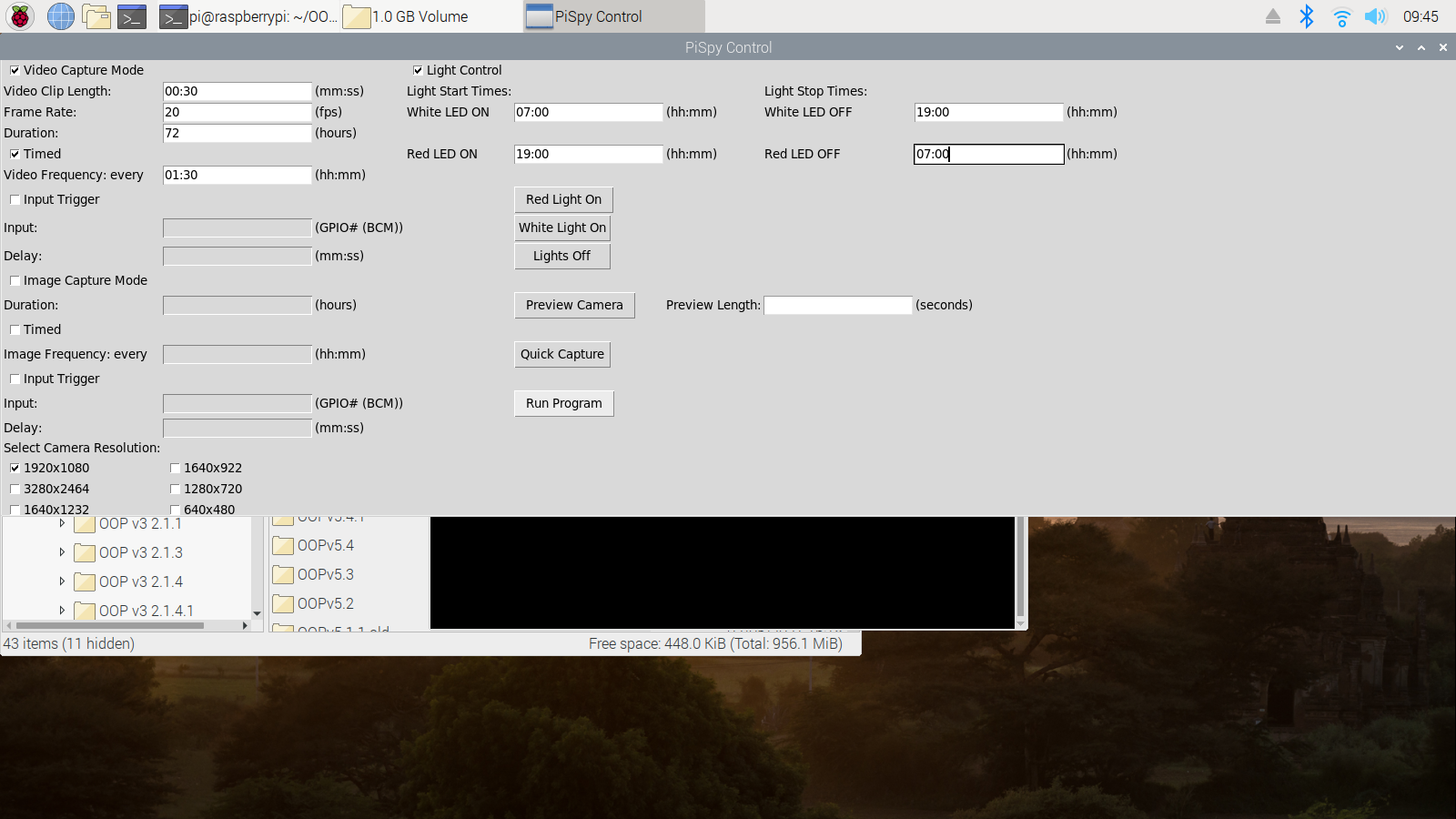

#### Input trigger

Input trigger mode can be used when the user wants an external source such as a motion sensor or force plate to determine when images or videos are recorded. See **Connecting GPIO sensors for input trigger** for more information on how to use different sensors for input trigger mode.

#### Light Control

If light control is desired, check the corresponding box and enter the times when each light should be turned on/off (hh:mm format, using a 24-hour clock). Using the red and/or white light buttons, turn on the light(s) you want on when the program starts running. In the image below, the white lights will be on from 7 AM to 7 PM, and the red lights will be on from 7 PM to 7 AM.

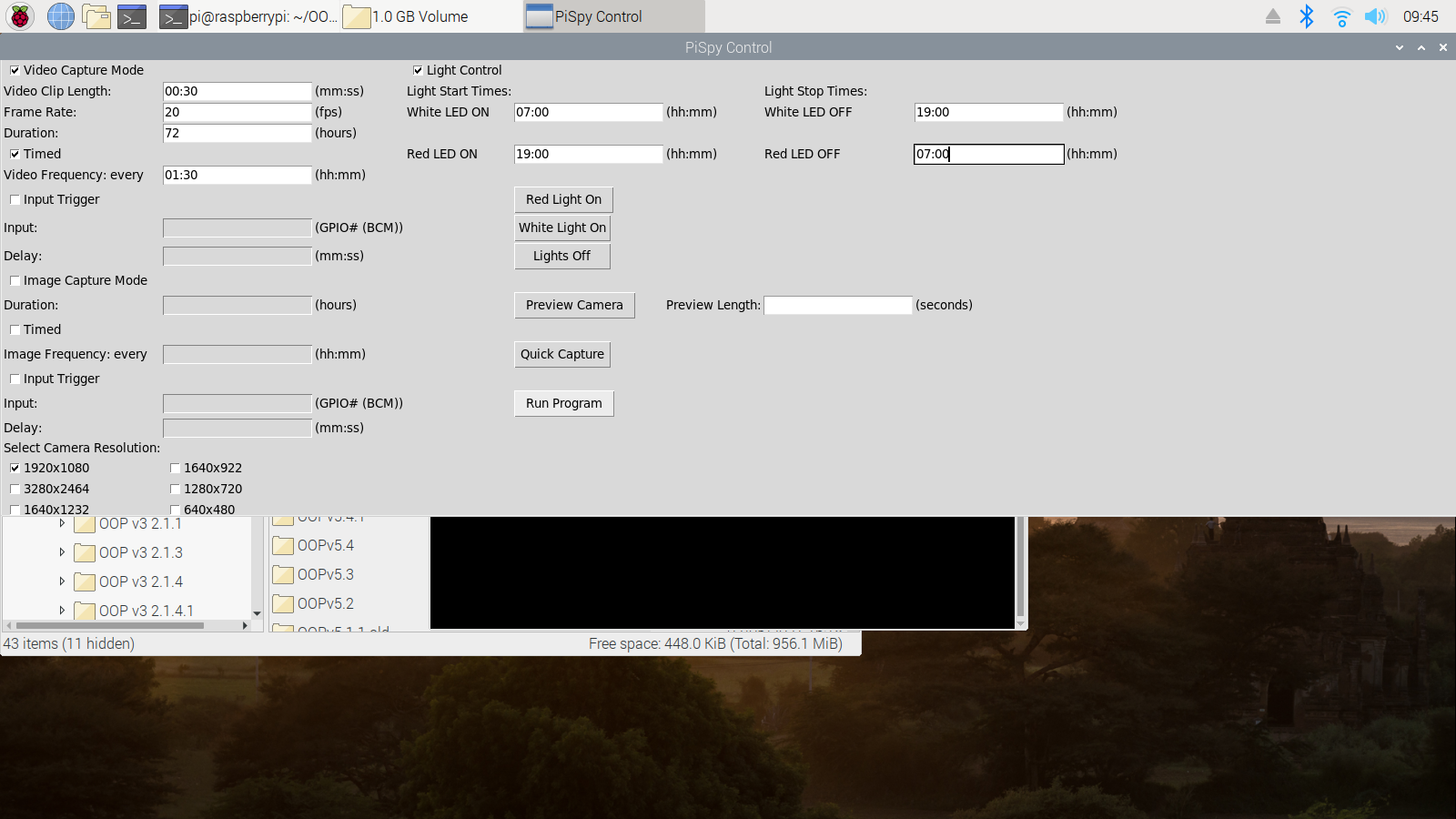

#### Note on Red Lights at Night

The default settings programmed into the PiSpy are what we found in testing to give the best quality night images/recordings. However, because of the PiCamera’s auto white balance settings, if the subject isn’t getting enough light or is sending back too much glare towards the camera, images/videos recorded at night can appear washed out. In addition to making changes to the PiSpy settings (see **Adjusting Default PiSpy Settings** below), changing the location and angle of the lights on the PiSpy, the proximity of the subject to the lens, and the height of the camera can all also help with improving night image quality.,

#### Quick Capture

If the user wants to manually take a picture or video, they can use the quick capture button. Select either video capture mode or image capture mode, and if taking a video also specify the clip length. Then, click the quick capture button to record.

#### Preview

To preview the output of the camera, the user can select the “preview’’ button. The preview length box allows the user to select how long the preview will last for (while the preview window is open, the user cannot interact with the PiSpy Control window). If no time is selected, the PiSpy will default to a 10 second preview.

#### Running the PiSpy Program

Once all the settings have been chosen, click the apply button to run the program.

Once the settings are applied, the PiSpy control window will close to run the program, and the chosen settings will be printed out to the terminal window. If the program needs to be stopped before the designated end time, perform a keyboard interrupt in terminal (control + c) or close the terminal window by clicking the X in the top right corner of the window. This will keep any previously recorded images/videos, but stop the program from recording any more, allowing the user to relaunch the control window and select new settings if desired.

### Connecting GPIO sensors for input trigger

#### Connecting a PIR motion sensor

A PIR motion sensor will have three pins: VCC, GND, and OUT. On some PIR motion sensors, the labels are under the cap; if this is the case, gently remove the cap to be able to identify the pins (the image below is a PIR sensor with the cap removed to show the pin labels). Using the image below as a reference (or by using the “pinout” command in terminal), connect the VCC pin to a 5V or 3.3V power supply pin on the Raspberry Pi, the GND pin to a ground pin on the Pi, and the OUT pin to any of the numbered GPIO pins on the Pi. GPIO board numbering differs between different models of the Raspberry Pi, so if you are not using the Raspberry Pi 3 you will need to consult a different chart when connecting the sensor. The OUT will output a voltage when the sensor is activated, which will be received by the Raspberry Pi and trigger the PiSpy to record an image/video (depending on selected settings). The diagram below is just a suggestion and does not need to be followed exactly; the sensor will work as long as GND is plugged into a ground pin, OUT to a GPIO pin, and VCC to a power supply.

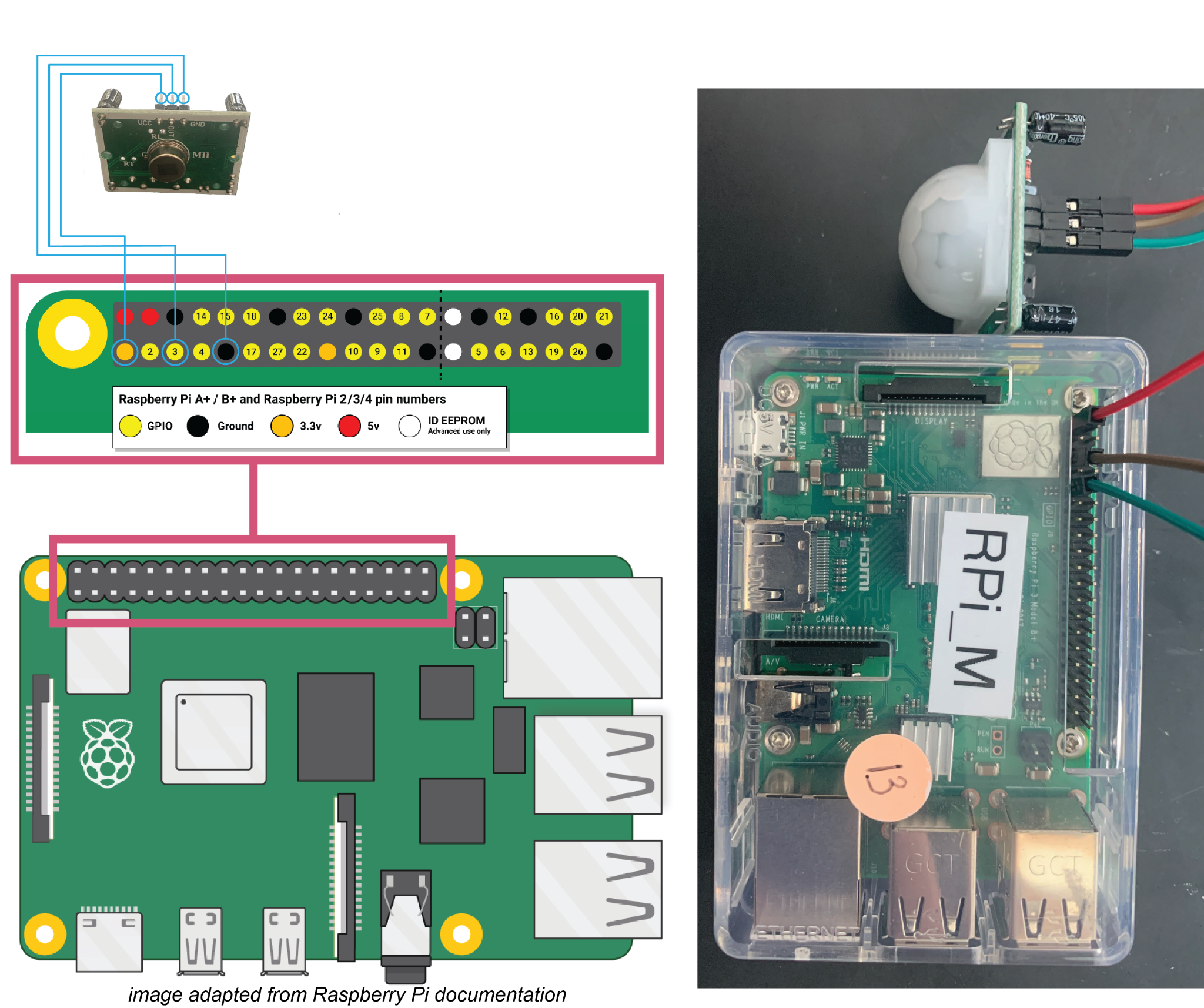

#### Connecting an IR break beam

An IR break beam has two components, an emitter and a receiver, but the wiring is similar to a motion sensor. The emitter emits a beam of IR light and has two wires, a red one that receives power from a 5V or 3.3V (the range will be better if using 5V) power supply on the Raspberry Pi, and a black one that connects to a ground pin. The receiver, which receives the signal from the emitter, has the same red and black wires which must be plugged into a different power and ground pin, but also has a white or yellow output wire that should be plugged into one of the general purpose GPIO pins. See the image below for reference. As noted above, the power can come from a 3.3V pin, but the range of the break beam will be smaller. Additionally, any GPIO pin can be used, not just 2 (just make sure to specify which one the input is plugged into when you run the PiSpy)

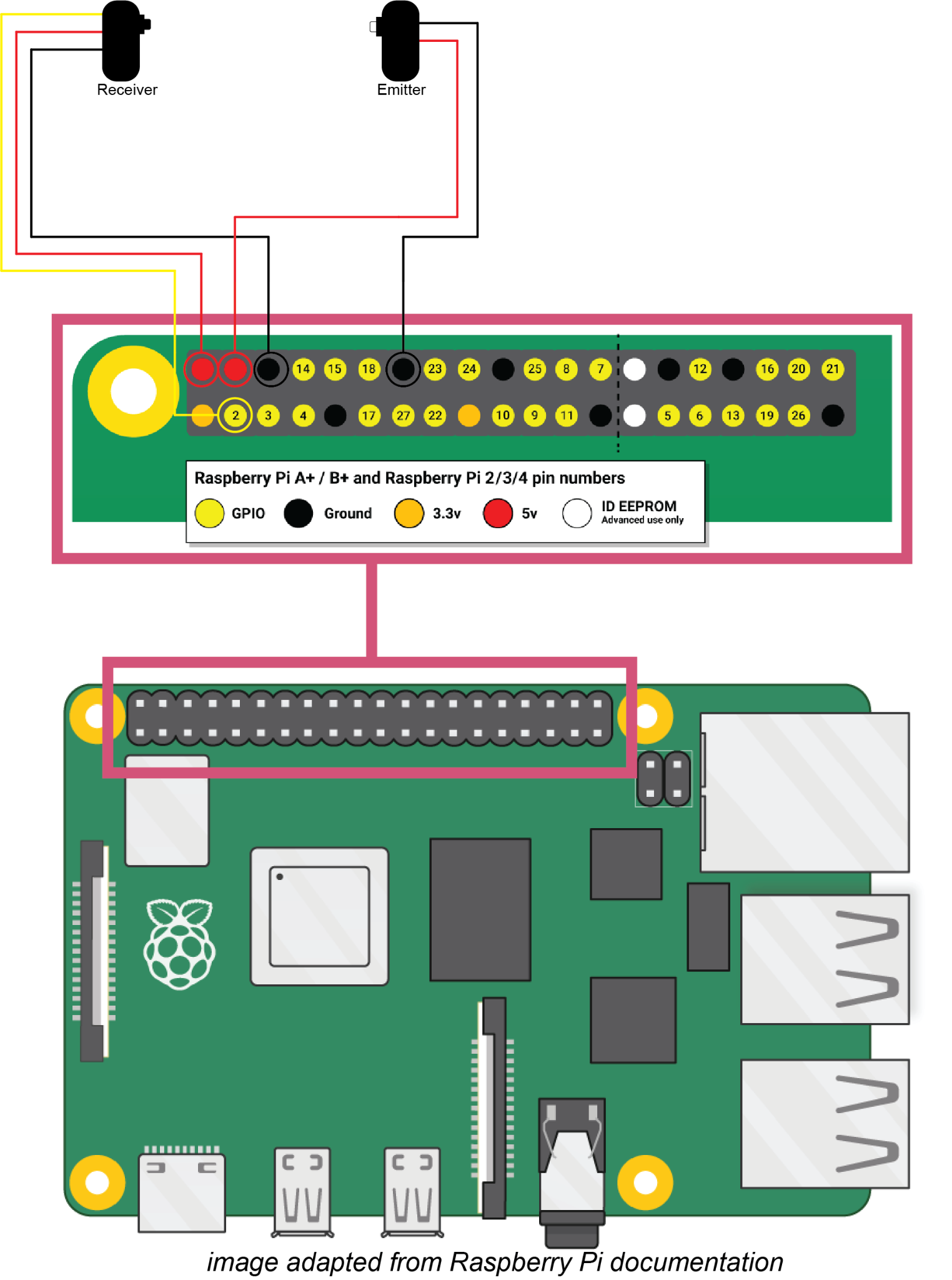

#### Note on PIR motion sensors

In our testing, several commercially available and inexpensive PIR motion simply do not function. If the motion sensor does not correctly trigger image/video acquisition, double check that every connection is in the proper position and secure, and if that doesn’t work, consider trying a different sensor.

#### Note on IR break beams

An IR break beam has two parts, an emitter that sends out IR light and a sensor that receives that signal and will signal to the computer when that beam is broken. In our testing, small objects (we were using ants) don’t always reliably break the beam and trigger the sensor. To make the sensor sensitive enough to trigger when an ant passed it, we used LEGOs® to decrease the total area that the sensor could receive IR signal from, and placed a single index card in front of the sensor as well. The index card is partially transparent to IR light, so when nothing else was between the sensor and the emitter the beam was not broken, but when an ant passed it would trigger video acquisition. Our suggestion is to test break beam sensitivity with your desired subject before running any experiment to confirm that the beam will trigger, and if it does not increase the sensitivity, which can be done with DIY tools and further testing as we did.

#### Note on BOARD vs BCM GPIO labeling

There are two different ways Raspberry Pi GPIO pins can be referred to. BOARD labeling refers to the actual position of the pin on the GPIO board, whereas BCM (Broadcom chip-specific pin numbers) refers to the lower-level functions of the pins. The PiSpy uses the BCM labelling throughout its code, so this is how the input source must be referred to when using input trigger mode.

#### Note on code for input triggers

Different input triggers function in slightly different ways, and the code (in the “Input_Trigger.py” file) must be slightly different for each of them. As currently deposited on GitHub, the code is written to work with an IR break beam, which has an output of 1 when the beam is not broken and 0 when the beam is broken. As advised in the Input_Trigger.py code, if using a PIR motion sensor, which outputs a 0 when it does not sense motion and a 1 when it does, change lines 56 and 113 of the Input_Trigger.py document from “if x == 0:” to “if i >= 1:” and comment out (place # signs before) lines 50-54 and 108-112. See the screenshots below for reference.

While we do not provide code/instructions for wiring of other types of input triggers besides PIR motion sensors and IR break beams, there are a considerable number of online resources available to help with any other types of sensor, many of which can be found directly on websites like Adafruit and raspberrypi.com when purchasing the sensors.

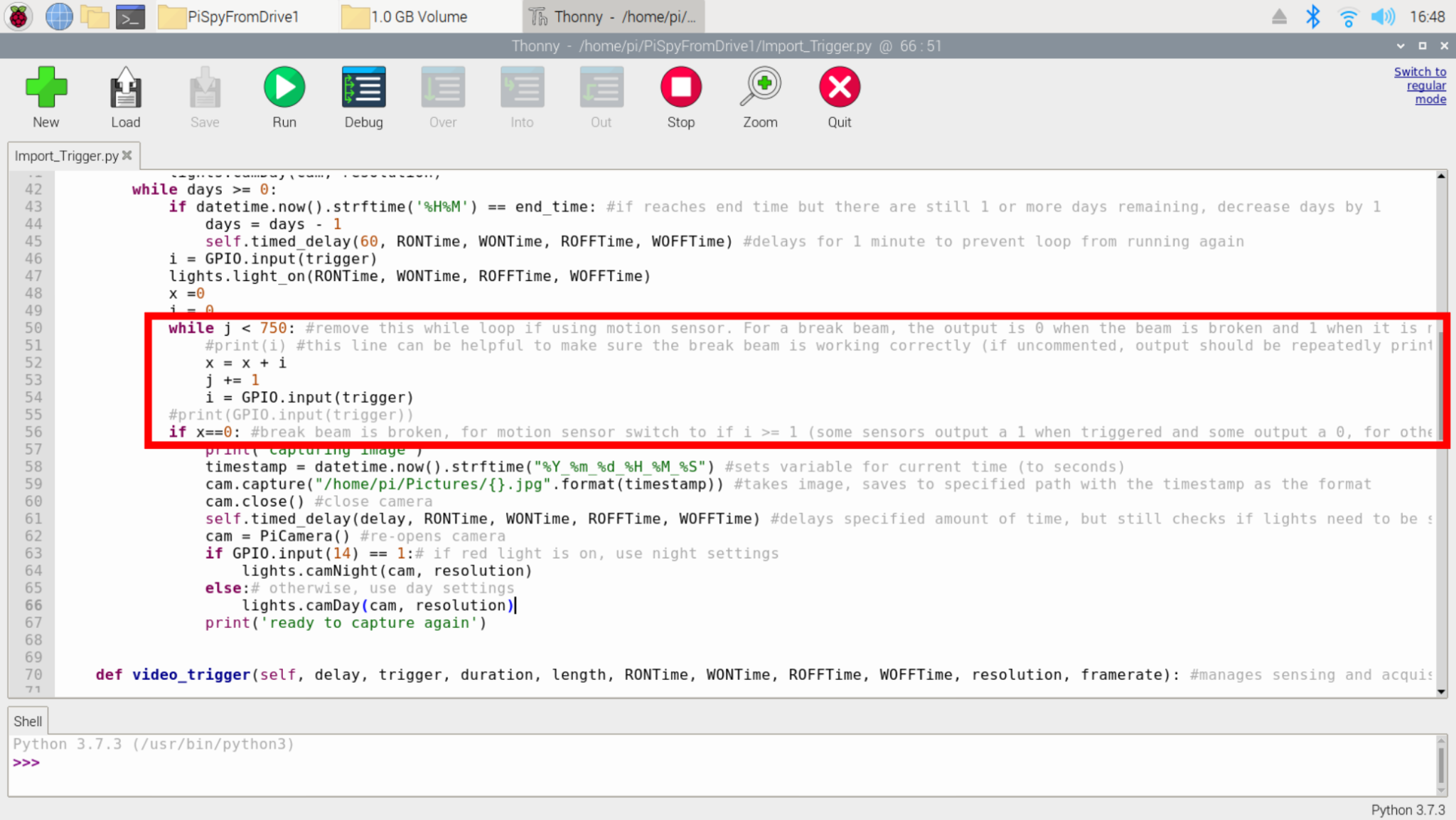

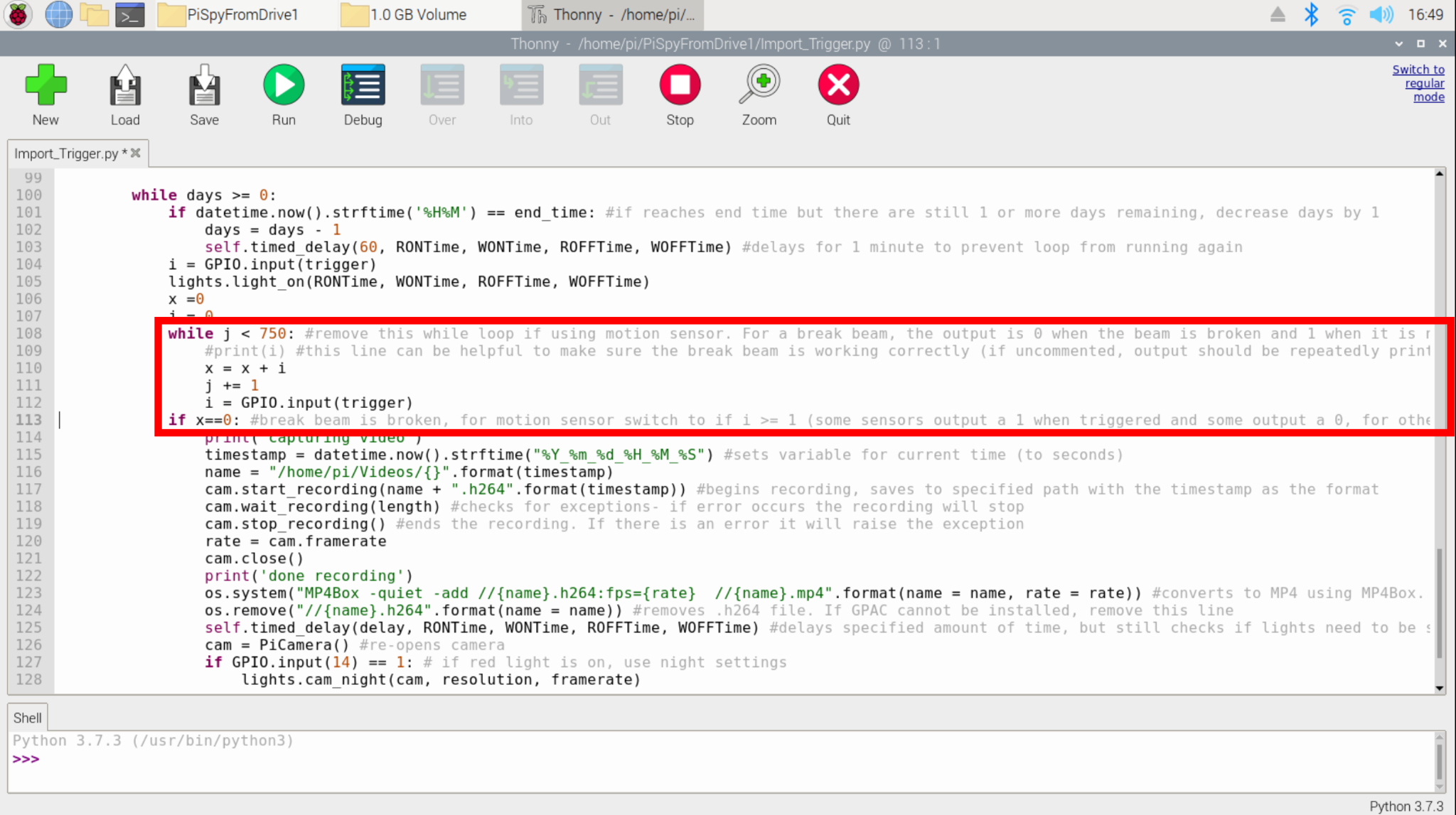

#### Using input trigger mode

In the PiSpy control window, specify which GPIO pin the trigger source is connected to, using the BCM format (the white text in the above diagram). Additionally, specify in the mm:ss format how long a delay is desired between image or video captures. Note that many sensors (i.e. PIR motion sensors) also have a mechanically adjusted delay setting, so this can also be adjusted to set the device’s delay. The PiSpy will use whichever delay is longer. In the image below, 1640x1232 images will be recorded when triggered by a sensor plugged into GPIO 4 with a 15 second delay. Recording will occur for 6 hours.

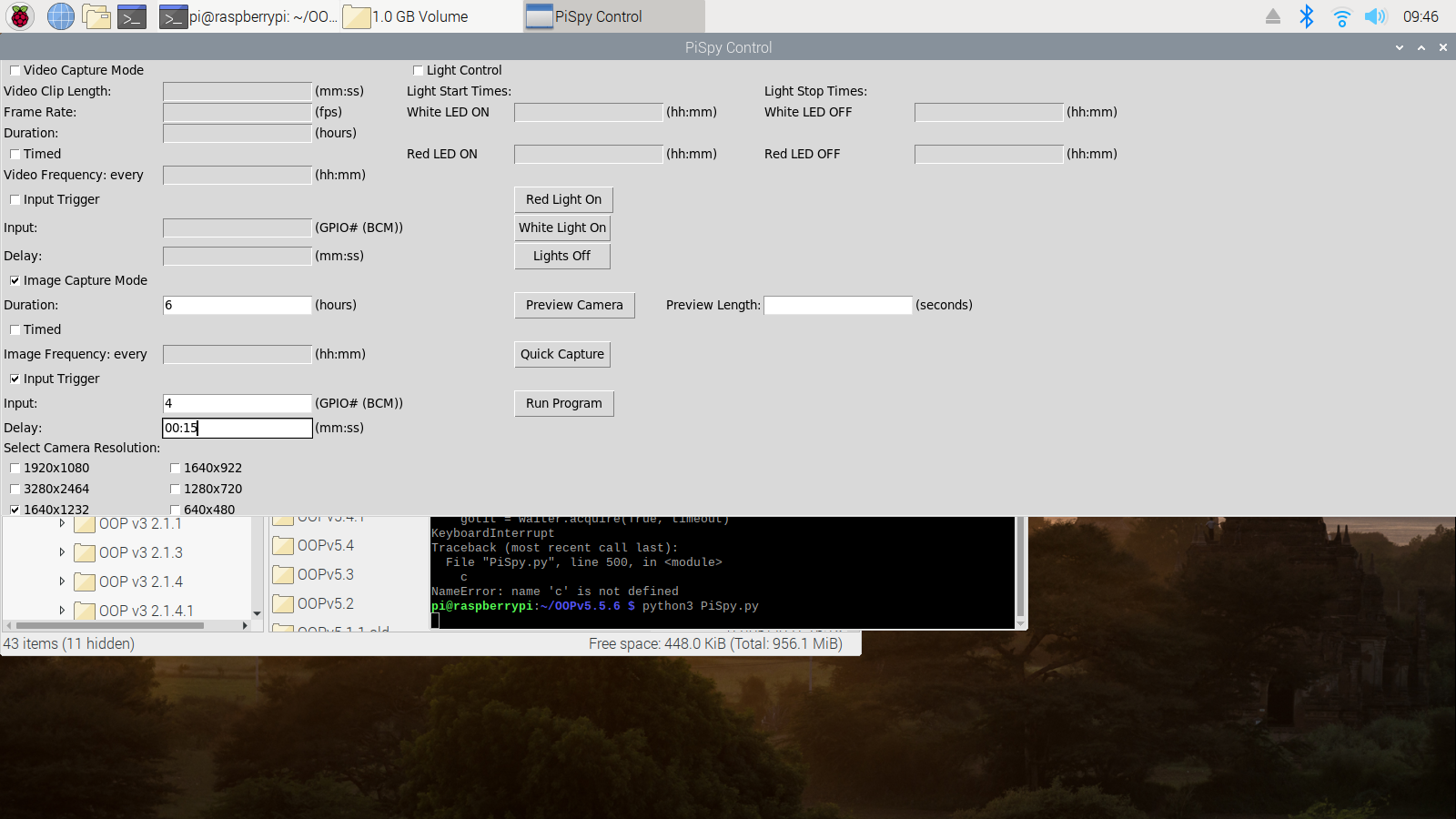

**Setting the time on a Raspberry Pi**

There are two ways to set the time on a Raspberry Pi. If the computer is connected to the internet and its time zone is set (which is done when the pi is first booted up, or can be done in the localisation tab of the Raspberry Pi Configuration window), the time will set automatically.

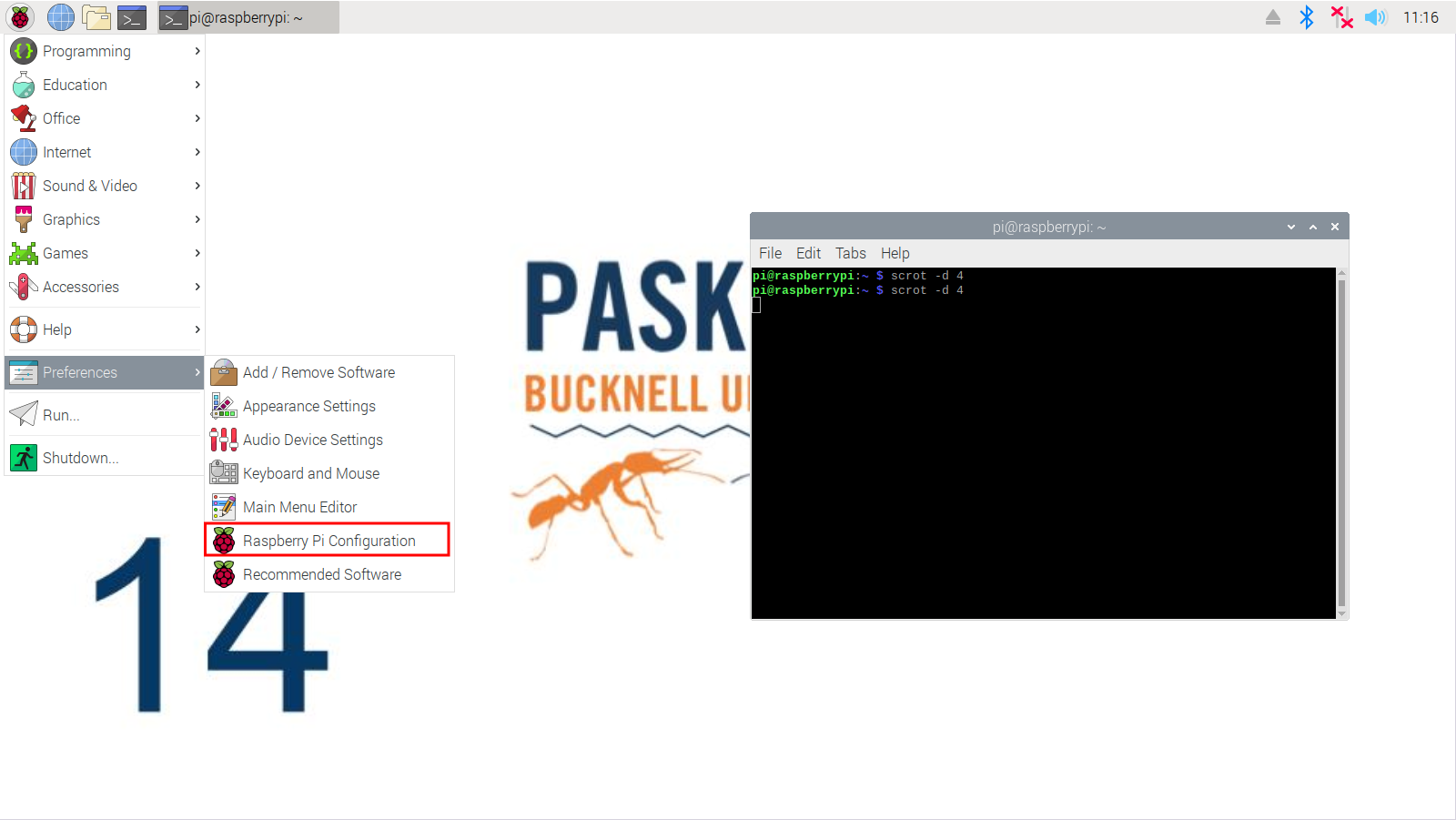

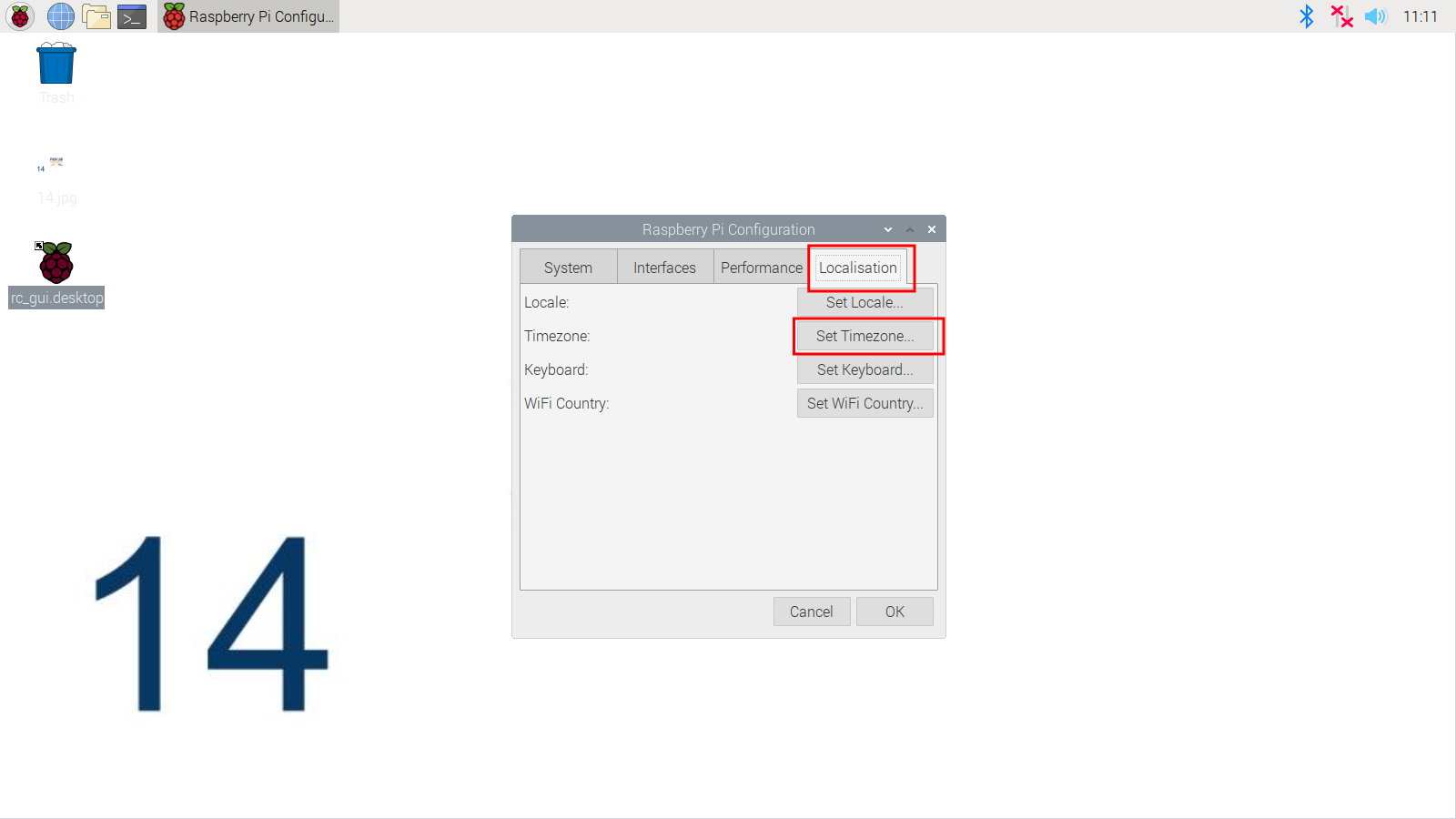

If the computer is not connected to the internet, the time will have to be set manually, and since Raspberry Pis do not have a built in clock this will have to be done every time it is powered back on. To manually set the time, open a terminal window and type the command (replacing the specifics with whatever the current date/time is):

*sudo date -s "Tue Jun 15 22:10:11 UTC 2021"*

### Adjusting Default PiSpy Settings

By default, the PiSpy camera will have an ISO of 200 during day recording (no lights on or white lights on), and an ISO of 800 during night recording (red lights on). At night, brightness (set to 43 on a scale from 1 to 100) and contrast (set to -8 on a scale from -100 to 100) are also lowered, as this helped improve image quality with the red LED lights on. If the user wants to adjust any of these settings, open the Day_Night_Mode.py file in the PiSpy folder. Then, make the desired adjustments under the camDay function (to change day settings) or the camNight function (to change night settings). Note that there are two camDay and two camNight functions (one for when framerate is specified and one for when it is not. To ensure a change in settings will always be implemented, make sure to adjust settings in both of these functions. In the screenshot below, the functions highlighted in red control day settings, and the ones in blue control night settings. Note, however, that image quality, especially at night, is also effected by the physical setup of the Pi. See *Note on Red Lights at Night* above for slightly more detail on this.

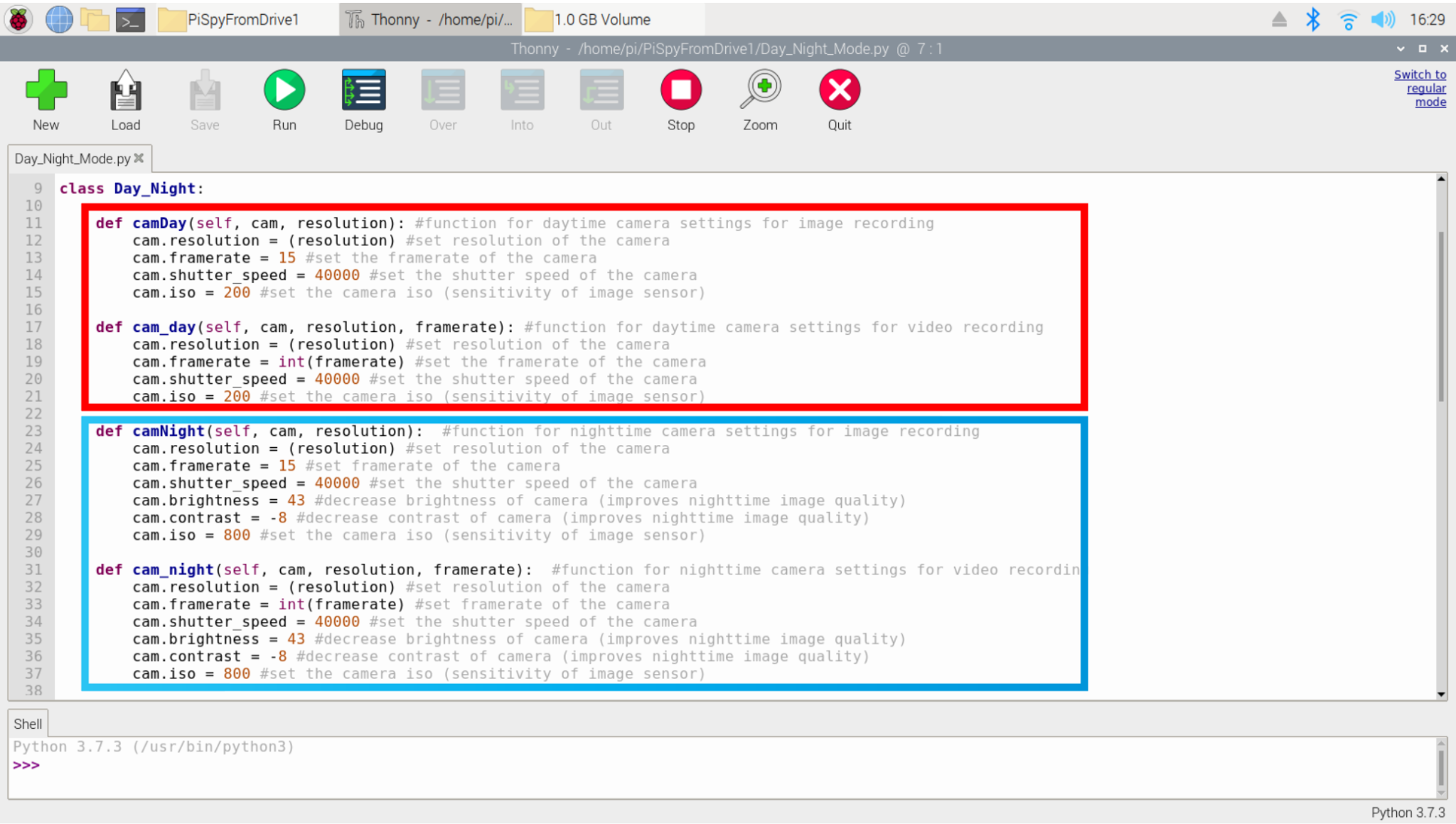

By default, the PiSpy will title images and videos with the time they were recorded (in the format year_month_day_hour:minute), and will store them to the default Pictures folder (/home/pi/Pictures).

To change either of these settings for image recording, open the “Image.py” file. To change the location the images are stored at, replace /home/pi/Pictures in line 20 with the path to the desired folder (make sure to leave the /{}.jpg after the path, as that is what specifies the file name). To change the file name, add any desired text before or after the {} in line 20 (the {} is a placeholder that tells Python where to put the timestamp in the filename). The format of the timestamp itself can be adjusted in line 18.

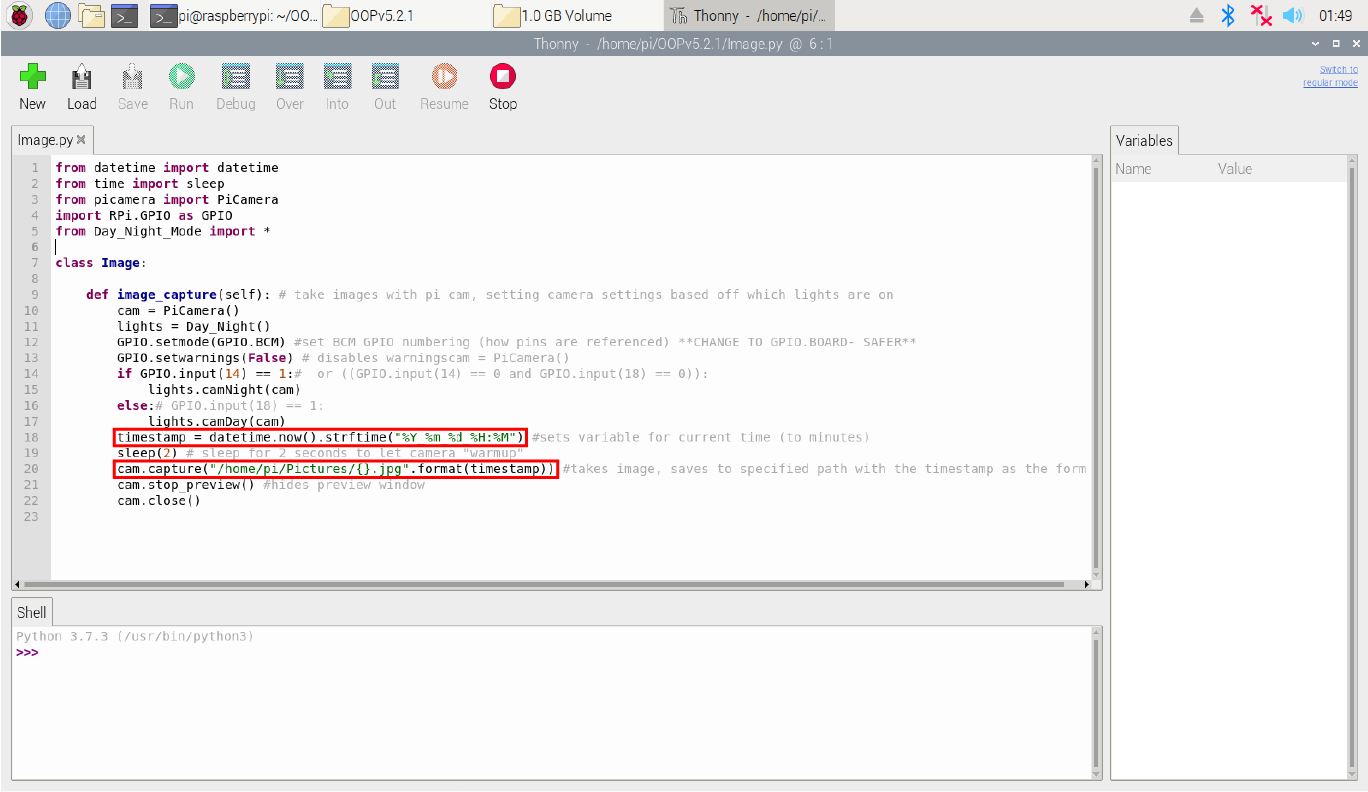

For video recording, open the “Record.py” file. The same steps as above can be used to adjust the file’s name and location, except the location/name is determined in line 24, and the format of the timestamp is determined in line 20.

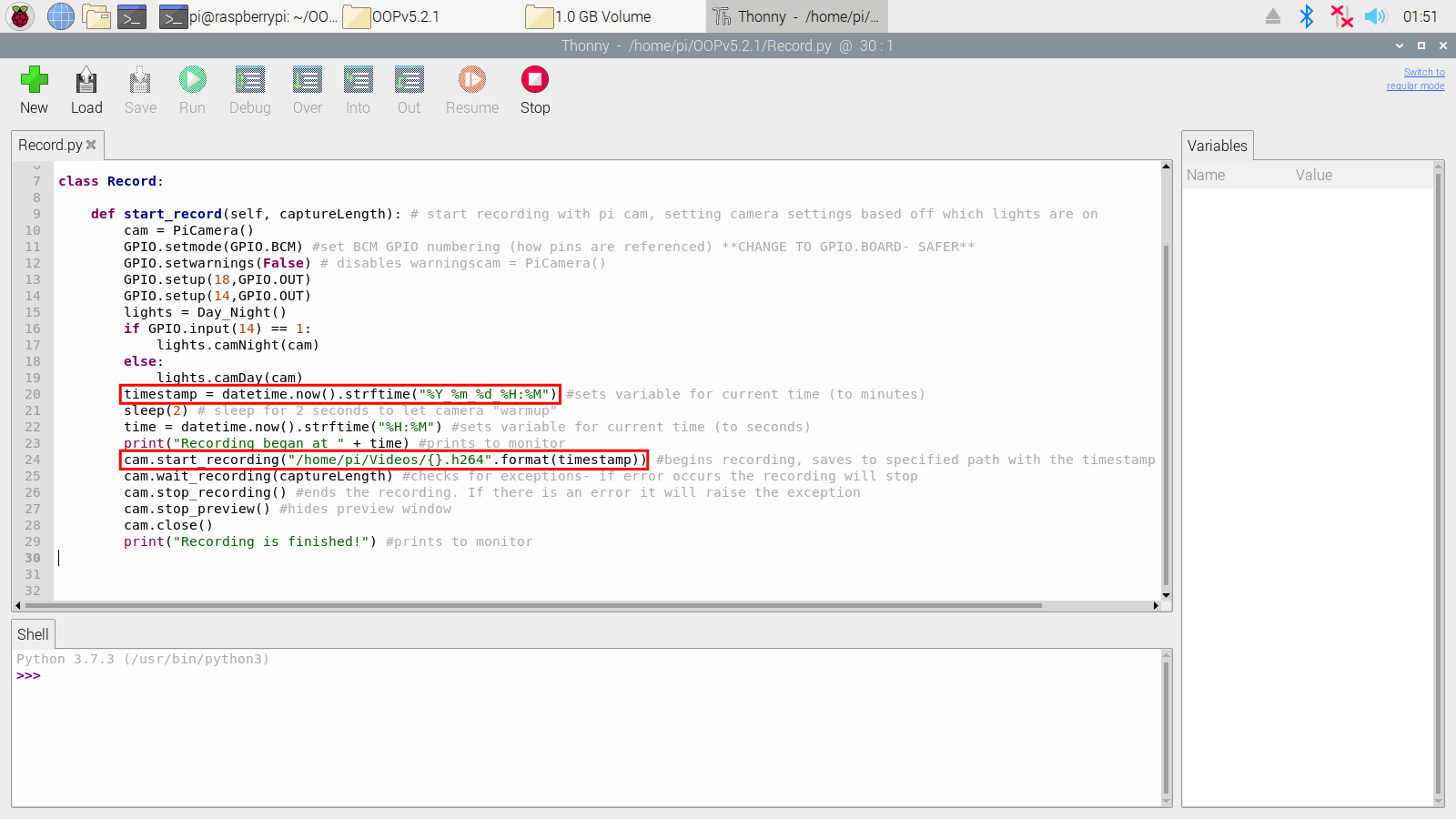

### LED PCB Lighting

Designs for white/red LED Printed Circuit Boards (PCBs) are on the PiSpy GitHub repository and can be used to purchase PCBs from fabrication businesses. One or more LED PCBs must be connected to the GPIO board on the Raspberry Pi for lighting control. When the Pi is off, connect the GPIO pins to the corresponding pins on the LED PCB(s) as shown in the diagram below. For GPIO positions, please refer to the pinout diagram above. Alternatively, the pinout command in terminal will display a diagram of the GPIO for your Raspberry Pi model.

If using other lighting outputs, please note the GPIO pins of interest and that the GUI code may need to be modified.

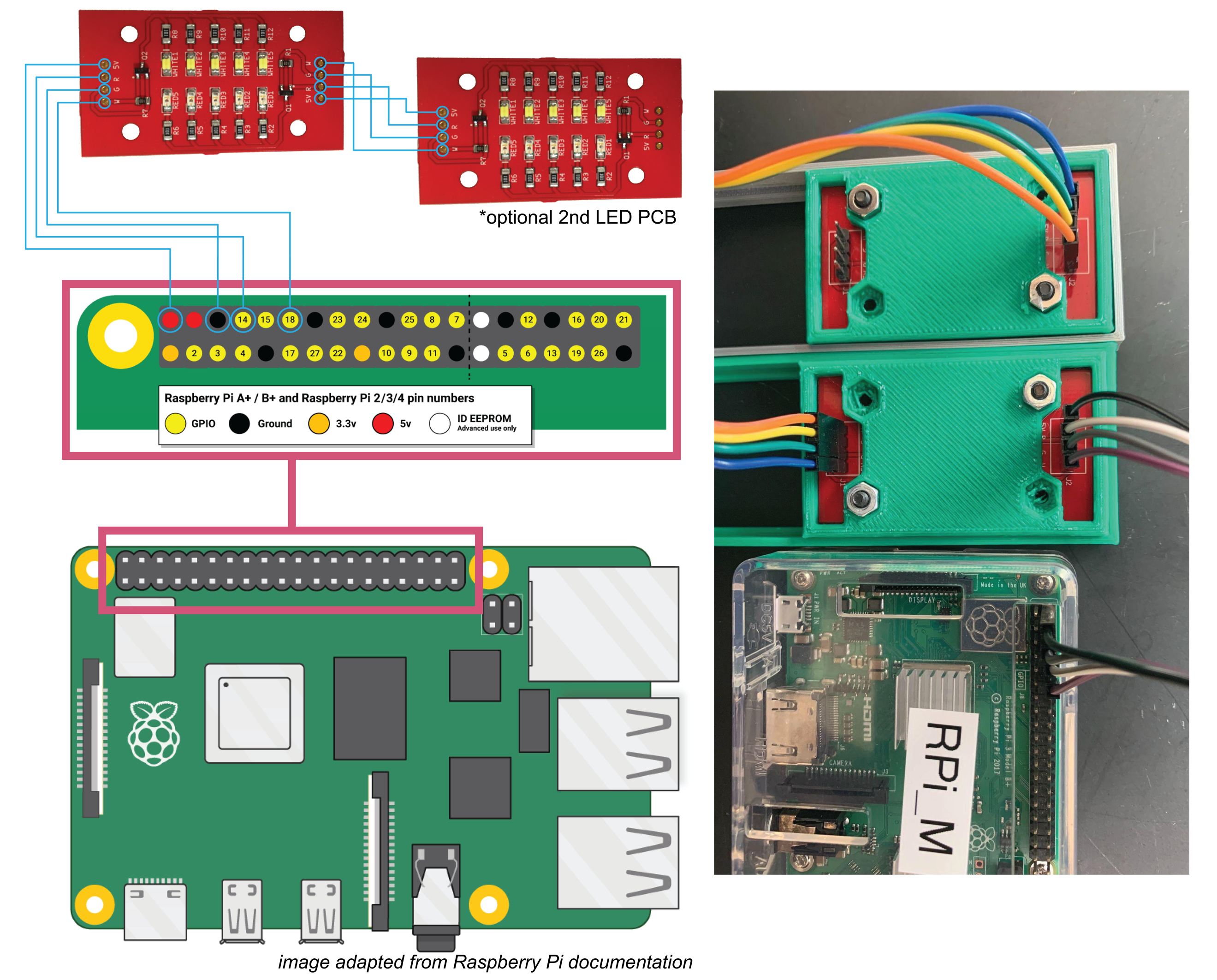

1. The legacy version of the Raspberry Pi OS must be used because the most recent OS update (2021.10.30), Bullseye, does not support the PiCamera python library used by the PiSpy.
